# Aberrant TSC-Rheb axis in Oxytocin receptor+ cells mediate stress-induced anxiety

**DOI:** 10.1101/2024.06.25.600464

**Authors:** Olivia Tabaka, Saheed Lawal, Rodrigo Del Rio Triana, Mian Hou, Alexandra Fraser, Andrew Gallagher, Karen San Agustin Ruiz, Maggie Marmarcz, Matthew Dickinson, Mauricio M. Oliveira, Eric Klann, Prerana Shrestha

**Affiliations:** Department of Neurobiology & Behavior, Stony Brook University, Stony Brook, NY 11794; Center for Neural Science, New York University, New York, NY 10003

## Abstract

Stress is a major risk for the onset of several maladaptive processes including pathological anxiety, a diffuse state of heightened apprehension over anticipated threats^1^. Pathological anxiety is prevalent in up to 59% of patients with Tuberous Sclerosis complex (TSC)^2^, a neurodevelopmental disorder (NDD) caused by loss-of-function mutations in genes for Tuberin (*Tsc2*) and/or Hamartin (*Tsc1*) that together comprise the eponymous protein complex. Here, we generated cell type-specific heterozygous knockout of *Tsc2* in cells expressing oxytocin receptor (OTRCs) to model pathological anxiety-like behaviors observed in TSC patient population. The stress of prolonged social isolation induces a sustained negative affective state that precipitates behavioral avoidance, often by aberrant oxytocin signaling in the limbic forebrain^3,4^. In response to social isolation, there were striking sex differences in stress susceptibility in conditional heterozygote mice when encountering situations of approach-avoidance conflict. Socially isolated male mutants exhibited behavioral avoidance in anxiogenic environments and sought more social interaction for buffering of stress. In contrast, female mutants developed resilience during social isolation and approached anxiogenic environments, while devaluing social interaction. Systemic and medial prefrontal cortex (mPFC)-specific inhibition of downstream effector of TSC, the integrated stress response (ISR), rescued behavioral approach toward anxiogenic environments and conspecifics in male and female mutant mice respectively. Further, we found that *Tsc2* deletion in OTRCs leads to OTR-signaling elicited network suppression, i.e., hypofrontality, in male mPFC, which is relieved by inhibiting the ISR. Our findings present evidence in support of a sexually dimorphic role of prefrontal OTRCs in regulating emotional responses in anxiogenic environments, which goes awry in TSC. Our work has broader implications for developing effective treatments for subtypes of anxiety disorders that are characterized by cell-autonomous ISR and prefrontal network suppression.

## Main

Social connection is a fundamental need for social animals including mice and humans – it encompasses affiliative care, social play, vocal communication, and allogrooming among conspecifics. Deprivation of social connection acts as a stressor that has variable effects on behavioral performance depending on duration of the experience, following the inverted U-shaped curve described by Yerkes and Dodson law^5^. Social isolation for a short period of time elicits social craving and increases social interaction as a way of restoring social homeostasis^6–9^. The allostatic load of prolonged social isolation, on the other hand, precipitates a negative affective state and excessive behavioral avoidance especially in genetically susceptible individuals^3,10^. The high prevalence of pathological anxiety in TSC suggests an abnormal emotional response to pervasive uncontrollable stressors. Although cognitive and social deficits, specifically in the domains of learning and memory, behavior flexibility, and social recognition, have been extensively studied in TSC mouse models^11–15^, there is a lack of understanding of neural correlates of TSC-associated anxiety at the multi-scale level of molecular substrates, neuromodulatory pathways, and neural circuits. Yet, TSC-associated neuropsychiatric disorders (TAND)^16–18^, including pathological anxiety, lead to the greatest burden of this multisystem disease^17,19,20^.

The hypothalamic neuropeptide, oxytocin (OT), is a crucial component for moderating the outcome of stress response^21–24^ and for conferring social salience^4^. The central effects of OT are mediated through its membrane-bound G-protein coupled receptors (OTRs) that are expressed in specific brain cells (OTRCs) in experience- and sex-dependent manner. At the network level, OT modulation has been shown to refine the salience of neuronal network activity, mainly by facilitating GABA-dependent increase in signal-to-noise ratio, although its role in glutamatergic and astrocyte modulation has been addressed recently^4,25,26^. OT is released from the paraventricular nucleus during a stressor and impedes the ensuing response of the hypothalamo-pituitary-adrenal (HPA) axis^27–29^. Midbrain OTRCs are implicated in social craving following acute social isolation^9^, whereas *Oxtr* gene expression in the central amygdala is downregulated during chronic social isolation leading to anxiety-related behaviors^30^. The medial prefrontal cortex (mPFC) integrates information from multiple brain regions including sensory, motor, and limbic systems, and exerts top-down control on emotion regulation in response to acute and chronic stressors. Although there is no direct evidence for a role of prefrontal OTRCs in social isolation, optogenetic silencing of OTRCs in adult mPFC leads to anxiety-like behaviors in a sex-dependent manner, mainly through their interaction with corticotropin release factor (CRF) stress-dependent signaling^31,32^. Chronic social isolation in adolescence reduces excitability of mPFC neurons in the deep layers that project to subcortical targets including the paraventricular thalamus^33^, and nucleus accumbens^34^, which results in impaired social recognition. On the other hand, chronic social isolation in adult male mice, but not juvenile mice, precipitates depressive- and anxiety-like behaviors^35^, indicating an age-dependent, domain-specific susceptibility to chronic social isolation^36^. Aside from its role in stress modulation, the mPFC is itself a target of the maladaptive stress response. Cellular and morphological changes, including dendritic atrophy and reduced myelination, occur in mPFC following chronic exposure to an emotional stressor^37–40^. Similarly, the OT system itself undergoes adaptive changes during chronic stress, for example by adjusting *Oxtr* gene expression, intrinsic properties of OT neurons or by modifying input signals to OTR neurons, leading to modulation of physiological set points to adapt to changing environments^4,41,42^. Boosting the function of the OT-OTR system has been effective in improving socioemotional behaviors in several cases of diseased brain states^43,44^.

To unravel the molecular and cellular correlates of TSC-associated pathological anxiety, we introduced functional inactivation of *Tsc2* in the OT system. *Tsc2* encodes a GTPase activating protein (GAP) for small G protein Rheb, which can associate with and activate both mammalian target of Rapamycin complex 1 (mTORC1)^45^ as well as PKR-like endoplasmic reticulum kinase (PERK)^46^.The latter effector triggers the ISR, a complex neuroprotective pathway aimed at reducing protein synthesis while allowing translation of select mRNAs to promote cell recovery and survival^47,48^. Prolonged ISR however is detrimental to neuronal function and limits plasticity mechanisms that build resilience in response to external psychological stressors^49,50^. We hypothesized that cell type-specific disruption of *Tsc2* in OTRCs might cause heightened susceptibility to social isolation and precipitate maladaptive anxiety-related behaviors by amplifying the signaling pathways negatively controlled by TSC complex. To test this hypothesis, we carried out heterozygous deletion of *Tsc2* in OTRCs and systematically examined the effects of prolonged social isolation on anxiety-related behaviors in male and female mice. Using pharmacological approaches, we determined PERK-ISR as the causal molecular mechanism that precipitates emotional susceptibility to social isolation in male mice. Further, we identified OTRCs in mPFC as the cellular substrates that mediate the behavioral and electrophysiological signatures of ISR-induced network suppression and stress-induced anxiety.

### Cell type-specific deletion of *Tsc2* in OTRCs alters energy balance during stress

To genetically target the OTRCs in mouse brain, we bred the OTR Cre BAC transgenic mice (founder line ON66)^51^ (**Fig. 1a, Extended Data Fig. 1a-i)** with *Tsc2* floxed mice^52^, which resulted in cell type-specific heterozygous deletion of *Tsc2* gene in OTRCs throughout development and adult life (**Fig. 1b**). Homozygous null mice were not viable. We then determined the changes in TSC2 protein abundance in mPFC OTR+ neurons that were labeled virally expressed double-floxed mCherry in adult wildtype (WT) and conditional heterozygous *Tsc2* (cHET) mice (**Fig. 1c)**. Heterozygous gene deletion of *Tsc2* resulted in an average 32% reduction in TSC2 protein level in OTR+ neurons. Interestingly, the cumulative relative distribution of TSC2 abundance in OTR+ neurons showed that approximately 60% of cells completely lack TSC2, reminiscent of the second-hit/ loss-of-heterozygosity model of TSC^53^, whereas a small percentage of OTR+ neurons have no change in the level of TSC2 compared to wildtype OTR+ cells (**Fig. 1d, e)**. We next assessed how *Tsc2* haploinsufficiency in OTRCs modifies the effects of prolonged social isolation. For this, mice were singly housed starting at 3 months of age and were fed *ad libitum* and provided with identical environmental enrichment, including bedding and nesting material, as when they were group housed **(Fig. 1f)**. Mice were weighed at 3 and 5 months of age, in group housed condition and after social isolation stress (SIS). We found that the bodyweight of wildtype males increased normally over two months of social isolation, from 3 to 5 months of age. However, similar weight gain was absent in the male cHET mutants (**Fig. 1g**). To assess whether this lack of weight gain was due to changes in feeding behavior, we restricted food access of socially isolated males for 24 hours and then measured the amount of chow consumed in the home cage over 10 min. There was no change in food consumption (**Extended Data Fig. 2a**), which indicated that the energy expenditure, which prolonged stress response can necessitate, is elevated in SIS-treated cHET males despite similar food intake. In the case of female mice, we found that cHET mice gained weight over the social isolation period however the WT females did not gain any weight (**Fig. 1h)**. Like males, both genotypes of female mice were comparable in food intake in their home cage post social isolation (**Extended Data Fig. 2b)**. These data indicate that genetic deletion of *Tsc2* in OTRCs renders opposite effects of stress susceptibility for maintenance of energy balance in male and female mice.

**Figure 1.**
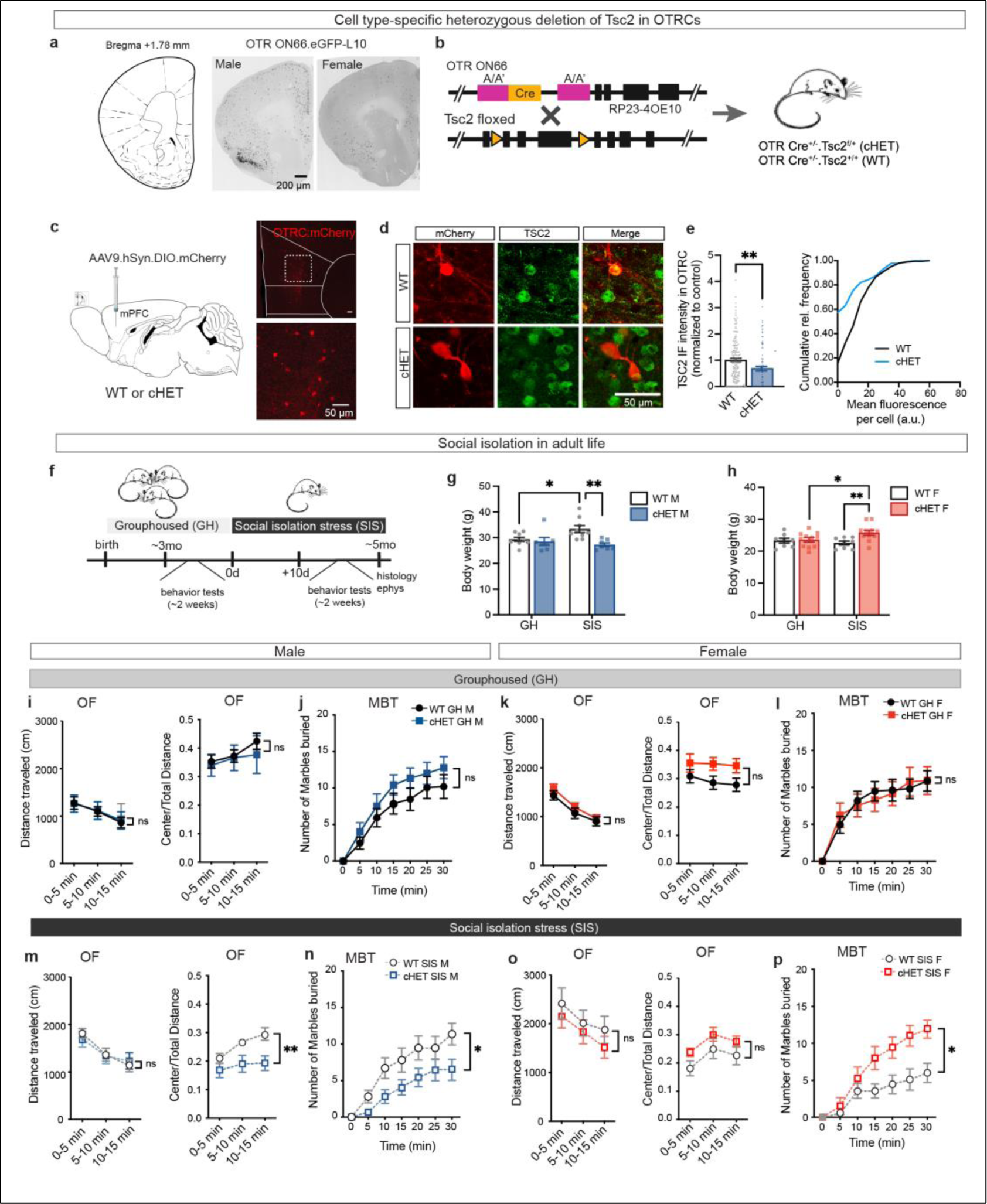
Cell-type specific heterozygous deletion of *Tsc2* in OTRCs. **a)** Cre recombinase activity in the prefrontal cortex of OTR Cre driver strain, founder line #ON66, bred to floxed GFP-tagged ribosomal protein L10. **b)** Breeding scheme for generating heterozygous deletion of *Tsc2* in OTRCs (OTR Cre^+/-^ .Tsc2^f/+^, cHET) and control Oxytocin Cre transgenic mice (OTR Cre^+/-^.Tsc2^+/+^, WT). **c)** Viral expression of floxed mCherry reporter in OTRCs within mPFC. **d)** Representative images of mPFC sections from WT and cHET mice, showing reduction of TSC2 protein expression in OTRCs within mPFC. Scale bar = 50 μm. **e)** Comparison of TSC2 immunofluorescence intensity in prefrontal OTRCs in WT and cHET mice (Left). Cumulative relative frequency of TSC2 immunofluorescence in WT and cHET mPFC. About 60% of OTRCs in cHETs have complete depletion of TSC2, whereas the remaining 40% OTRCs have variable reduction of TSC2 (right). **f)** Schematic for behavioral timeline for measuring the effects of social isolation stress (SIS) in adult life. **g)** Disruption of energy balance in cHET males: bar graph showing natural gain in bodyweight over 2 months of SIS in WT males but lack of such a gain in cHET male mice. **h)** Bar graph showing loss of body weight gain in WT female mice upon social isolation, but normal weight gain in cHET female mice over the period of SIS indicating stress resilience. **i)** Normal exploratory behavior (left) and thigmotaxis (right) in grouphoused male WT and cHET mice. **j)** Normal marble burying behavior in grouphoused male mice of both genotypes. **k)** Normal spontaneous locomotion and thigmotaxis in grouphoused female mice, and **l)** normal marble burying behavior in grouphoused females. **m)** SIS exposure has similar effects on exploratory behavior in WT and cHET males (left) but there is a significantly greater stress susceptibility in cHET males for thigmotaxis (right). **n)** Marble burying behavior is significantly reduced in socially isolated male cHET mice compared to WT mice. **o)** Spontaneous locomotion (left) and thigmotaxis (right) are comparable in SIS-exposed WT and cHET female mice. **p)** Socially isolated female WT mice bury fewer marbles compared to cHET mice. Statistical tests: e) Student’s unpaired t-test; g, h) Two-way ANOVA with Bonferroni post-hoc test; i-p) RM Two-way ANOVA with Bonferroni post-hoc test, *p<0.05, **p<0.01, ns not significant. Sample size: e) n = 189 WT cells from 4 mice, and 89 cHET cells from 3 mice; g-p) n = 8-18 mice/group.

### Sex differences in stress susceptibility in anxiety-like behaviors

To determine if there were sex differences in the effect of SIS on behavior, we longitudinally tested adult mice in a battery of anxiety-related tests when they were group-housed and after they were socially isolated for 10 days. We found strikingly different behavioral effects of *Tsc2* gene deletion and SIS on male and female mice. While there was no remarkable effect of genotype on anxiety-like behaviors (**Fig. 1i-l)**, social isolation caused a small increase in the spontaneous locomotion in male mice and led to avoidance of the center zone in the open field arena indicating an increase in thigmotaxis (**Extended Data Fig. 3a, b**). This increase in anxiety-like behavior in the open field was significantly amplified in cHET male mice (**Fig. 1m)**. Specifically, while the SIS-exposed WT males progressively adapted to the open field and ventured into the center towards the later phases of the trial, such an adaptation was lacking in SIS-exposed cHET males. In the elevated plus maze test, we observed a trend-level difference in the percent duration, but not in percent entries, in the open arm for SIS-exposed cHET mice compared to the grouphoused condition (**Extended Data Fig. 3c).** In the marble burying test, we found that SIS blunted the motivation in cHET males to bury marbles (**Fig. 1n)**, while causing no significant effect on the wildtype males (**Extended Data Fig. 3d, e)**. Because the socially isolated cHET males had no phenotype in self-grooming (**Extended Data Fig. 3f)** and rearing (**Extended Data Fig. 3g)**, we deduced that their stereotypic behaviors are not affected by the gene deletion and that the decrease in marble-burying behavior is due to a lack of motivational drive in performing an ethologically relevant behavior. In the case of female mice, there were no remarkable difference in open field behavior in WT and cHET female mice (**Fig. 1o**). SIS increased spontaneous locomotion in the open field and decreased exploration of the center zone in the initial phase (0-5 min) of the trial for both WT and cHETs (**Extended Data Fig. 4a, b**). However, social isolation was anxiolytic for the female cHETs in the elevated plus maze, where they spent more time exploring the open arm than when they were group-housed (**Extended Data Fig. 4c**). In marble burying test, we found that social isolation reduced motivational drive in female WT mice to bury marbles **(Extended Data Fig. 4d)**. In contrast, the female cHET mice were buffered from the effects of SIS on marble burying behavior (**Fig. 1p)**, even though there was a short but insignificant delay in these mice to match the behavior of the grouphoused counterparts (**Extended Data Fig. 4e**). The stability in marble-burying behavior in stress-exposed cHET females were not caused by compensatory stereotyped repetitive tendency since these mice were comparable to WT female mice in self-grooming (**Extended Data Fig. 4f**) and rearing (**Extended Data Fig. 4g**). These findings indicate that reduced *Tsc2* gene dosage in OTRCs have opposite effects on males and females when presented with an approach-avoidance conflict imposed by anxiogenic spaces.

To distinguish whether the anxiety displayed by socially isolated male mutants is a form of enduring trait-anxiety that transcends different situations or is situation-directed state-anxiety, we next tested mice in the three-chamber social interaction (3CSI) assay where they make a choice between interacting with a sex-matched novel conspecific or an inanimate object. In the group housed condition, both male and female cHET mice showed strong social preference for a sex-matched conspecific (**Fig. 2a, b**). Upon social isolation, we found that the male cHETs, like other male groups, continued to prefer the male stranger mice compared to the object, demonstrating a robust social preference index (**Fig. 2c, d**). Probed further in a dyadic social interaction test in their home cage, socially isolated male cHETs displayed a notable increase in social investigation, including sniffing and allogrooming, without any change in antagonistic attacks (**Fig. 2e-g**). In addition, this enhanced dyadic social investigation did not extend to courtship rituals toward sexually receptive female conspecifics (**Extended Data Fig. 5a-d**). We deduced that socially isolated male cHETs seek social interaction for buffering stress caused by anxiogenic environments. In sharp contrast, we found that socially isolated female cHETs, that were stress-resilient in anxiogenic environments, lacked a preference for the stranger female conspecific over an inanimate object (**Fig. 2i, j**). Social interaction is likened to a fundamental need and deprivation of social interaction for a finite duration leads to seeking of social feedback, like food-seeking behavior of animals in the hunger brain-state. Absence of seeking of social interaction in female cHETs after social isolation indicates a new set-point for these animals where they are habituated to asocial disposition, thus a lack of social motivation. In other words, prolonged social isolation produced behavioral adaptations in female cHETs to cope with anxiogenic environments but at the same time abolished the value of social interaction. An alternative possibility is that prolonged social isolation stress caused social anxiety, a negative affective state, in female cHETs however this is less likely given their stress resilience in anxiogenic environments.

**Figure 2.**
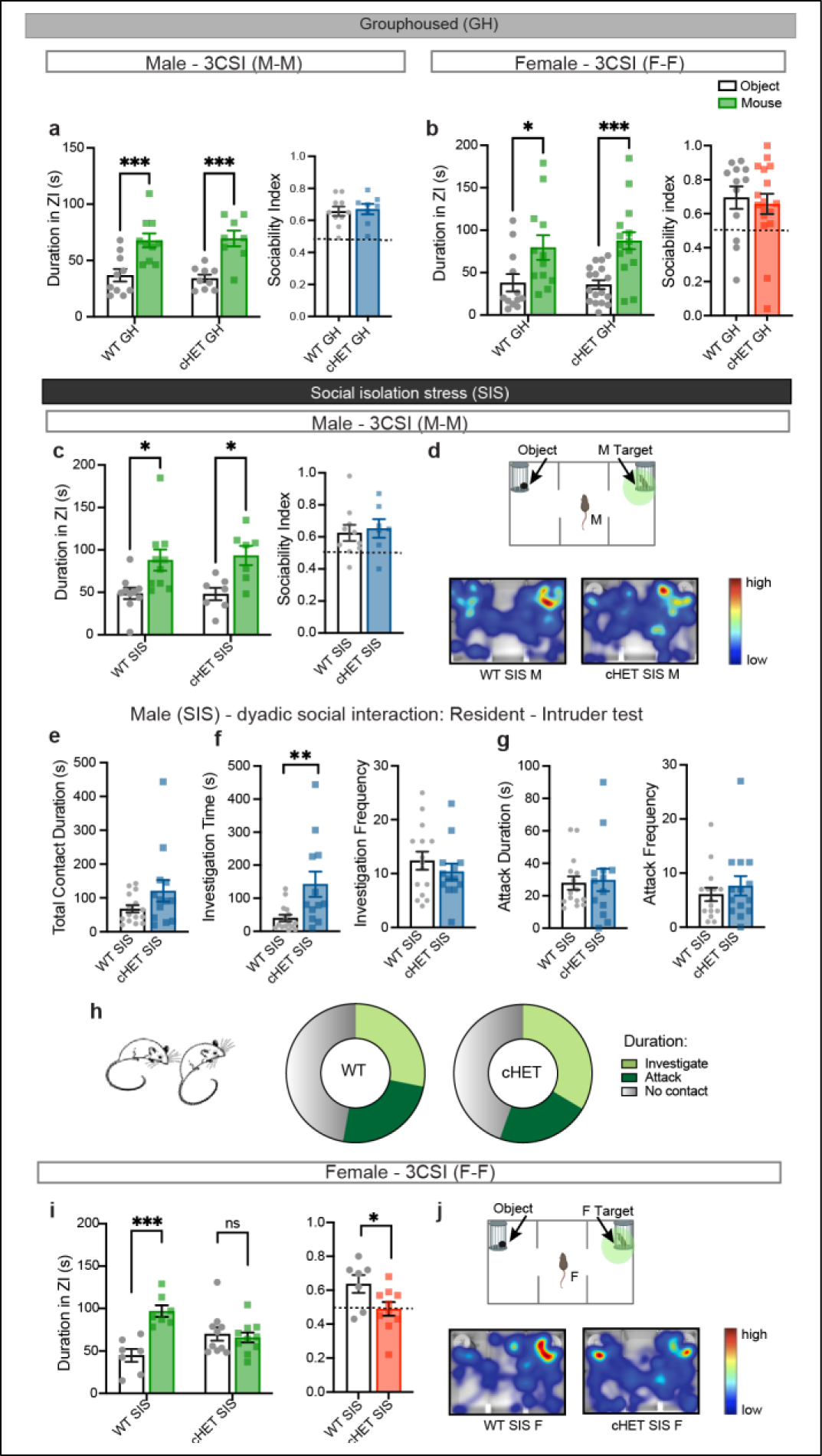
Sex differences in social interaction of cHET mice. **a)** In a 3-chamber social interaction (3CSI) test, grouphoused male mice displayed social preference and spent more time in the zone of interaction with the stranger male compared with the object. **b)** Grouphoused female mice also displayed robust social preference toward sex-matched conspecific. **c)** SIS-exposed male mice continue to have robust social preference toward male stranger in the 3CSI test. **d)** Behavior schematic for 3CSI, and representative heat map for social interaction. **e)** No significant difference in total contact duration is observed between SIS-exposed WT and cHET male mice in the resident-intruder test. **f)** SIS-exposed male mutants spend more time investigating, i.e, sniffing and grooming, the stranger male compared to WT mice, and spend more time during each investigation. **g)** SIS-exposed mice have normal aggressive bouts as wildtype mice. **h)** Venn diagram showing relative time spent in investigation and attack. **i)** Socially-isolated female mutants show no preference for the female stranger over an object, unlike all other groups including group-housed female mutants that prefer the social stimulus. Sociability index is reduced for SIS-exposed female cHET mice. **j)** Behavior schematic for 3CSI, and representative heat map for social interaction. Statistical tests: a, b, c, i) Left -Two-way ANOVA with Bonferroni post-hoc test; Right – Unpaired t-test; e-g) Unpaired t-test. *p<0.05, **p<0.01, ****p<0.001, ns not significant. Sample size: n = 7 – 15 mice/ group.

### Selective anxiolytic effects of pharmacological inhibition of downstream TSC effectors

Having established that social isolation produced a sustained anxiety state selectively in the male cHET mice, we sought to restore normal affective behaviors in anxiogenic environments. For this purpose, we measured the changes in key TSC-controlled signaling pathways in OTRCs deficient in TSC2. The GTPase activating protein (GAP) domain in TSC2 has GAP activity toward the small G protein Ras homolog enriched in brain (Rheb). Rheb has been shown to directly activate mammalian target of rapamycin complex I (mTORC1) and PKR-like endoplasmic reticulum kinase (PERK) in parallel^46,54^. Rheb is predominantly localized in the perinuclear area in the cytoplasm, and transient interaction of farnesylated Rheb is sufficient to activate mTORC1^55^, which is primarily localized in the lysosomes. Rheb enhances the binding of substrates, such as eukaryotic initiation factor 4E binding proteins (4E-BPs) and ribosomal protein S6 kinase (S6K1) to mTORC1, leading to an increase in cap-dependent translation, ribosome biogenesis, and cell growth. On the other hand, independent of mTORC1, Rheb also directly interacts with PERK, an ER-associated stress sensor protein, and triggers the integrated stress response via PERK-mediated phosphorylation of eukaryotic initiation factor 2α (eIF2α). Integrated stress response leads to a shutdown of general translation. Rheb activation can thus lead to opposing pathways that control nascent protein synthesis, and under physiological conditions, TSC complex has a critical role in negatively regulating Rheb. We thus sought to determine the effects on mTORC1 and PERK effectors in Tsc2-depleted OTRCs, specifically phosphorylation of ribosomal protein S6 (S6RP, Ser235/236) and eIF2α (Ser51). We injected a viral vector expressing double-floxed mCherry into the mPFC of wildtype or cHET mice to identify OTRCs and subjected the mice to social isolation stress followed by immunohistochemistry. We found that deletion of *Tsc2* from OTRCs results in upregulation of both mTORC1 and PERK pathways, as evident by increase in phosphorylation of S6RP (70%) and eIF2α (67%) (**Fig. 3a-c)**. These data show that rather than acting as a binary switch, TSC-Rheb axis exerts parallel control over the mTORC1 and PERK pathways (**Fig. 3d**). Further, this suggests that individual or combinatorial targeting of the downstream effectors of TSC, including both mTORC1 and PERK, could be effective in restoring normal behavioral function in the mutant mice.

**Figure 3.**
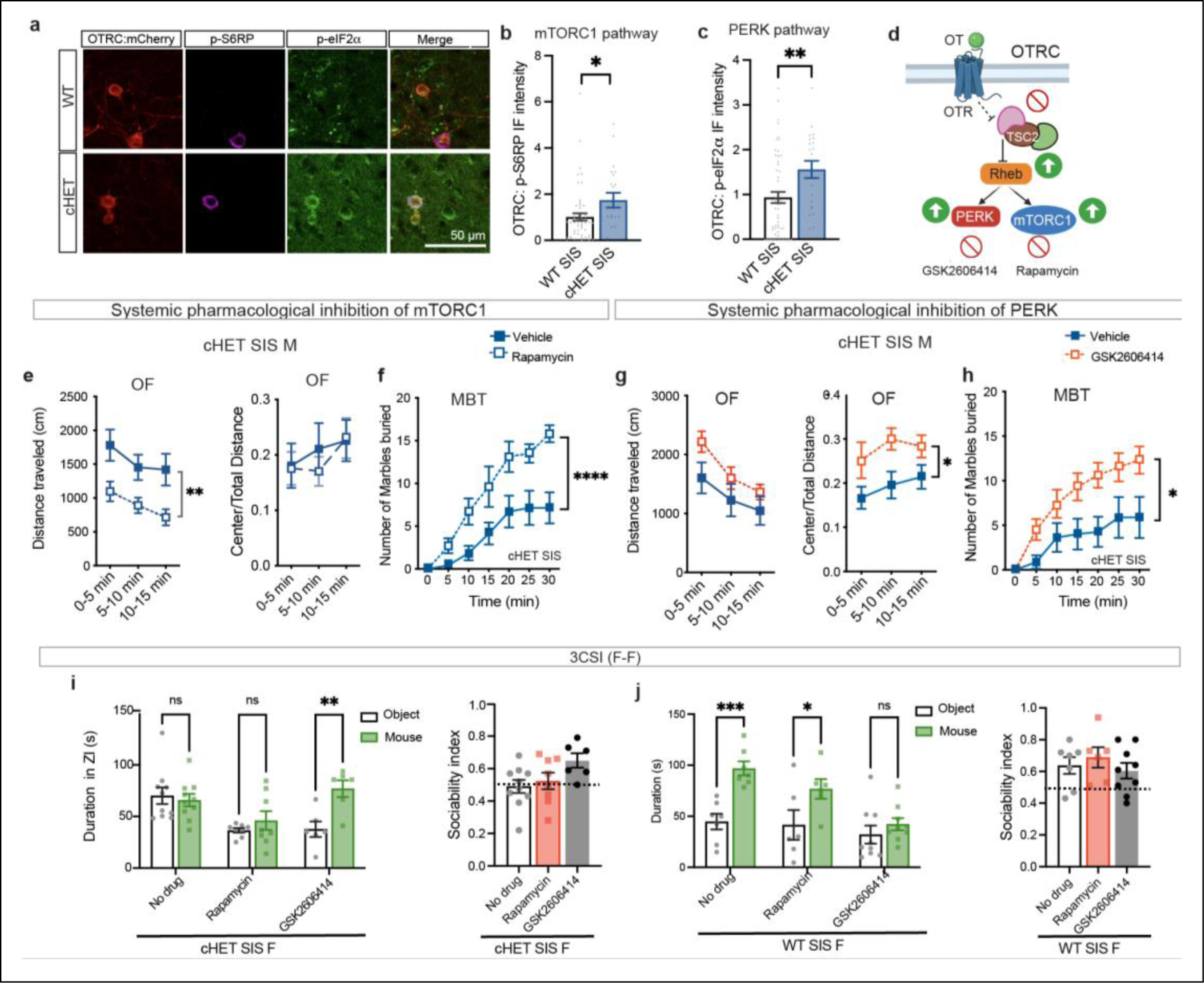
Systemic inhibition of mTORC1 and PERK pathways. **a)** Representative image showing immunohistochemistry for phosphorylated S6 (p-S6RP) and p-eIF2α. **b)** Optical density of p-S6RP immunofluorescence intensity, a proxy readout for mTORC1 pathway, in OTRCs is significantly increased in cHET mPFC compared to WT mPFC. **c)** Optical density of p-eIF2α IF intensity, a proxy readout for PERK pathway, is similarly increased in cHET mPFC. **d)** Molecular signaling schematic showing how TSC2 disruption in OTRCs affects downstream effectors. **e)** Subchronic systemic administration of Rapamycin, a specific mTORC1 inhibitor, lowers spontaneous locomotion of cHET males in the open field arena, but does not change thigmotaxis behavior. **f)** Rapamycin rescues the motivational drive of singly housed cHET mice to bury marbles in the marble burying test. **g)** Systemic inhibition of PERK with GSK2606414 does not alter exploratory behavior in SIS-exposed cHET males but rescues approach toward center zone in the open field. **h)** PERK inhibition also rescues marble burying behavior in singly-housed cHETs. **i)** PERK inhibition, but not Rapamycin, leads to effective rescue of social preference in singly housed cHET female mice. **j)** Rapamycin does not change social approach whereas PERK inhibition in SIS-exposed WT females abolishes social preference, indicating PERK-mediated modulation of social approach. Statistical tests: f-k) RM Two-way ANOVA with Bonferroni post-hoc test; l-o) Two-way ANOVA with Bonferroni post-hoc test. *p<0.05, **p<0.01. ***p<0.001. b-c) n=54 cells from WT mice, and 21 cells from cHET mice (3 mice/group). e-j) n = 6-14 mice/group

Prior studies have shown that blocking mTORC1 signaling in adult *Tsc2* global heterozygote mice can rescue deficits in the cognitive and social domains^11,56^. Acute PERK inhibition, although not tested in TSC mouse models, has been shown to improve cognitive function by boosting translation output^57,58^. On the other hand, chronic PERK inhibition is associated with impaired behavioral flexibility and working memory deficit^59,60^. Hence, we asked whether subchronic inhibition of either mTORC1 or PERK could rescue stress-precipitated anxiety in Tsc2 cHET males and social avoidance in cHET females. For this, we systemically administered a specific inhibitor of mTORC1, rapamycin (5 mg/kg *i.p.*), or vehicle for 3 days to WT and cHET mice that were socially isolated. The drug regimen for rapamycin was effective in significantly reducing phosphorylation of S6RP (Ser240/244) by about 33%, indicating inhibition of mTORC1 (**Extended Data Fig. 6a, b**). We proceeded with testing drug-treated cHET and WT males in anxiogenic environments of open field and marble burying arenas. Rapamycin, while effective at reducing stress-induced increase in open field activity, did not correct thigmotaxis in male mutants (**Fig. 3e**). The same dose of Rapamycin also corrected stress-induced hyperactivity in wildtype males (**Extended Data Fig. 6c**), suggesting that this effect was genotype-independent. In the marble burying test, Rapamycin restored marble burying behavior in socially isolated cHET males (**Fig. 3f**) while having no consequence in socially isolated WT male mice (**Extended Fig. 6d**). On the other hand, subchronic inhibition of PERK had no effect on spontaneous locomotion in open field, but it restored normal approach toward the center of the arena in cHET mutants (**Fig. 3g**). PERK inhibition in WT males further increased open field activity but did not alter thigmotaxis (**Extended Data Fig. 6e**). In the marble burying test, GSK2606414 rescued the marble burying behavior in cHET males (**Fig. 3h**), while further elevating this behavior and eliciting stereotypy in WT males (**Extended Data Fig. 6f**). Overall, rapamycin and PERK inhibition had complementary effects on exploratory behavior (**Extended Data Fig. 7a, b**), but both restored the grouphoused-like marble burying behavior in cHET males (**Extended Data Fig. 7c, d**). We also found that social preference in female cHET mice was selectively rescued by PERK inhibition but not by rapamycin treatment (**Fig. 3i**). Of note, while rapamycin did not have any negative effects on social preference in WT females, PERK inhibition eliminated their social approach (**Fig. 3j**). These results indicate that PERK activity is closely titrated during social interaction – deviation in either direction leads to maladaptive social avoidance. All together, despite the activation of both mTORC1 and PERK arms downstream of TSC-Rheb, our findings show that the primary pathway responsible for aberrant emotional and social behaviors in heterozygous mutant males and females is the PERK-mediated ISR.

### Dysfunctional OTRCs in mPFC mediate stress-induced behavioral avoidance

To determine the locus of network disruption that precipitates stress-induced behavioral avoidance in TSC, we targeted prefrontal OTRCs for genetic manipulation in mutant males and females. We bilaterally injected a lentivirus expressing double-floxed shRNA for Rheb along with an AAV expressing double-floxed mCherry to both identify OTRCs and mediate knockdown of Rheb (**Fig. 4a**). With this strategy, Rheb shRNA expression in mPFC OTRCs targeted the Rheb mRNA for degradation and reduced the protein abundance of Rheb by 23.70% in wildtype mice and by 46.55% in the cHET mice compared to mCherry controls (**Fig. 4b-d).** We next tested whether knocking down Rheb in OTRCs restores behavioral approach-avoidance balance in socially isolated mutant mice. We found that mPFC-restricted Rheb knockdown increased both open field activity and center zone approach in cHET male mice, mimicking the effects of systemic PERK inhibition (**Fig. 4e, Extended Data Fig. 8c**). Importantly, this genetic manipulation also restored time-dependent adaptation in center approach in the open field. Rheb knockdown in WT prefrontal OTRCs led to an increase in spontaneous activity but no change in thigmotaxis (**Extended Data Fig. 8a**). In the marble burying test, targeted Rheb knockdown further reduced the marble burying behavior in cHET males whereas it did not alter this behavior in WT male mice (**Fig. 4f**). On the other hand, genetic reduction of Rheb in prefrontal OTRCs rescued social approach in socially isolated female cHET mice but did not alter social preference in wildtype controls (**Fig. 4g**). These data indicate that TSC-Rheb axis in mPFC OTRCs largely contributes to stress-induced behavioral avoidance observed in male and female cHETs. By the process of elimination, we deduced that OTRCs in mPFC are not involved in generating the motivational drive in male cHETs to bury marbles. The lack of motivation in performing an ethologically relevant behavior is likely a result of OTRC malfunction in other brain regions outside of mPFC, such as the striatal circuits where earlier studies have implicated TSC complex in marble burying behavior^61^.

**Figure 4.**
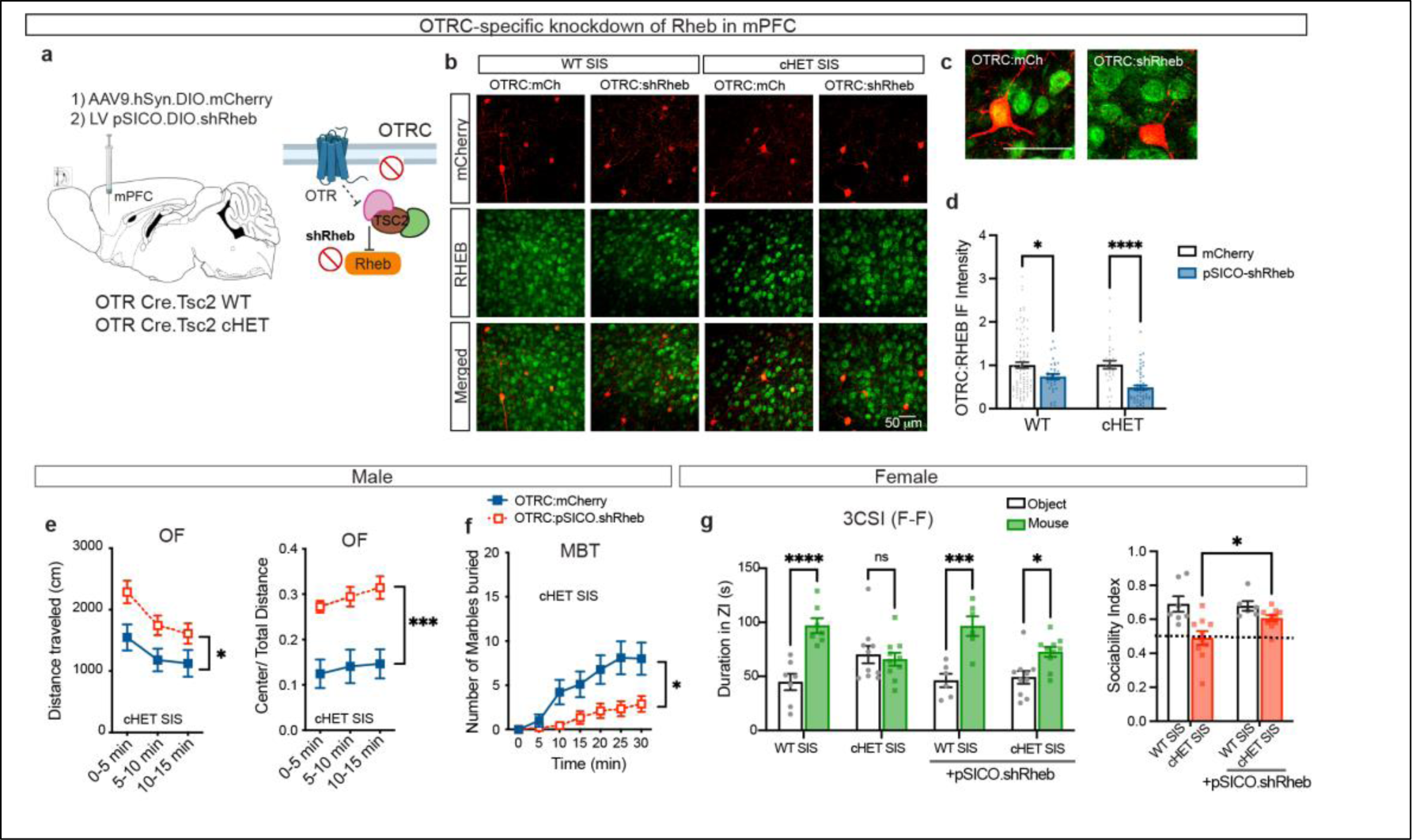
Cell type-specific knockdown of Rheb in prefrontal OTRCs. **a)** Schematic showing viral expression of mCherry and pSICO shRNA for Rheb in mPFC OTRCs. **b)** Representative images from immunohistochemistry carried out with anti-Rheb and mCherry. **c)** Representative inset images showing knockdown of RHEB. **d)** Optical density of RHEB immunofluorescence in OTRCs is significantly decreased upon shRheb expression in both WT and cHET mice. **d)** OTRC-specific Rheb knockdown increases spontaneous locomotion in SIS-exposed cHET male mice and **e)** increases approach toward the center zone in open field. **f)** Rheb knockdown in OTRCs decreases the motivational drive in cHET males to bury marbles. **g)** OTRC Rheb knockdown rescues social preference in cHET female mice but does not alter social approach in WT females. Statistical tests: c, g-h) Two-way ANOVA with Bonferroni post-hoc test; d-f) RM Two-way ANOVA with Bonferroni post-hoc test. *p<0.05, **p<0.01. ***p<0.001, ****p<0.0001. Sample size: c) n=33-99 cells/group from 3 mice/ group; d-h) n=6-10 mice/ group.

Given that PERK-ISR pathway in OTRCs is the primary contributor to the molecular aberrations precipitating stress susceptibility, we predicted that protein homeostasis is altered in these cells. To determine whether protein synthesis is disrupted in socially isolated cHET mice, we labeled OTRCs in mPFC using the viral approach to express mCherry, subjected mice to social isolation, and then performed fluorescent non-canonical amino acid tagging (FUNCAT)^62^ to label nascent peptides in mPFC slices for 2h *ex vivo* (**Fig. 5a**). Combining click chemistry with immunohistochemistry for mCherry, we found that there was an average 28% reduction in protein synthesis in cHET OTRCs compared to WT OTRCs within the mPFC (**Fig. 5b, c**). Analysis of cumulative relative frequency further showed that 30% of OTRCs in cHET mPFC have undetectable nascent protein synthesis (**Fig. 5d**). Considering the importance of the translation machinery in supporting synaptic plasticity, these data indicate that mPFC OTRCs function sub-optimally in modulating the network excitability. To determine whether *Tsc2* deletion in OTRCs affects the network activity in mPFC, we recorded evoked orthodromic synaptic potentials in socially isolated WT and cHET mice. It has previously been shown that majority of prefrontal OTRCs targeted using the ON66 Cre strain are GABAergic inhibitory neurons (80%) that express Somatostatin (SST, 80%)^31,51^. We stimulated superficial layer (L2/3) and recorded field excitatory postsynaptic potential (fEPSP) from inner (L5) cortical layer in mPFC slices and bath-applied a highly specific OTR agonist, [Thr4, Gly7]-oxytocin (TGOT) 20 min after the baseline recording (**Fig. 5e, Extended Data Fig. 9a**). After 10 min of TGOT treatment, we continued to record fEPSPs for 30 min. Although TGOT stimulation elicited a significant increase in fEPSP amplitude in WT male mPFC, cHET mPFC exhibited a deficiency in output spiking activity of inner cortical layers (**Fig. 5f, g**). Interestingly, in the case of female WT mPFC, TGOT stimulation did not produce network excitation and further, there was no difference in TGOT response between the female cHET and WT mPFC (**Extended Data Fig. 9b, c**). Li et al^31^ previously showed that the inhibitory postsynaptic current (IPSC) elicited by OTR interneurons in L5 pyramidal neurons is larger in females. This may explain the lack of network excitation with TGOT stimulation in WT females.

**Figure 5.**
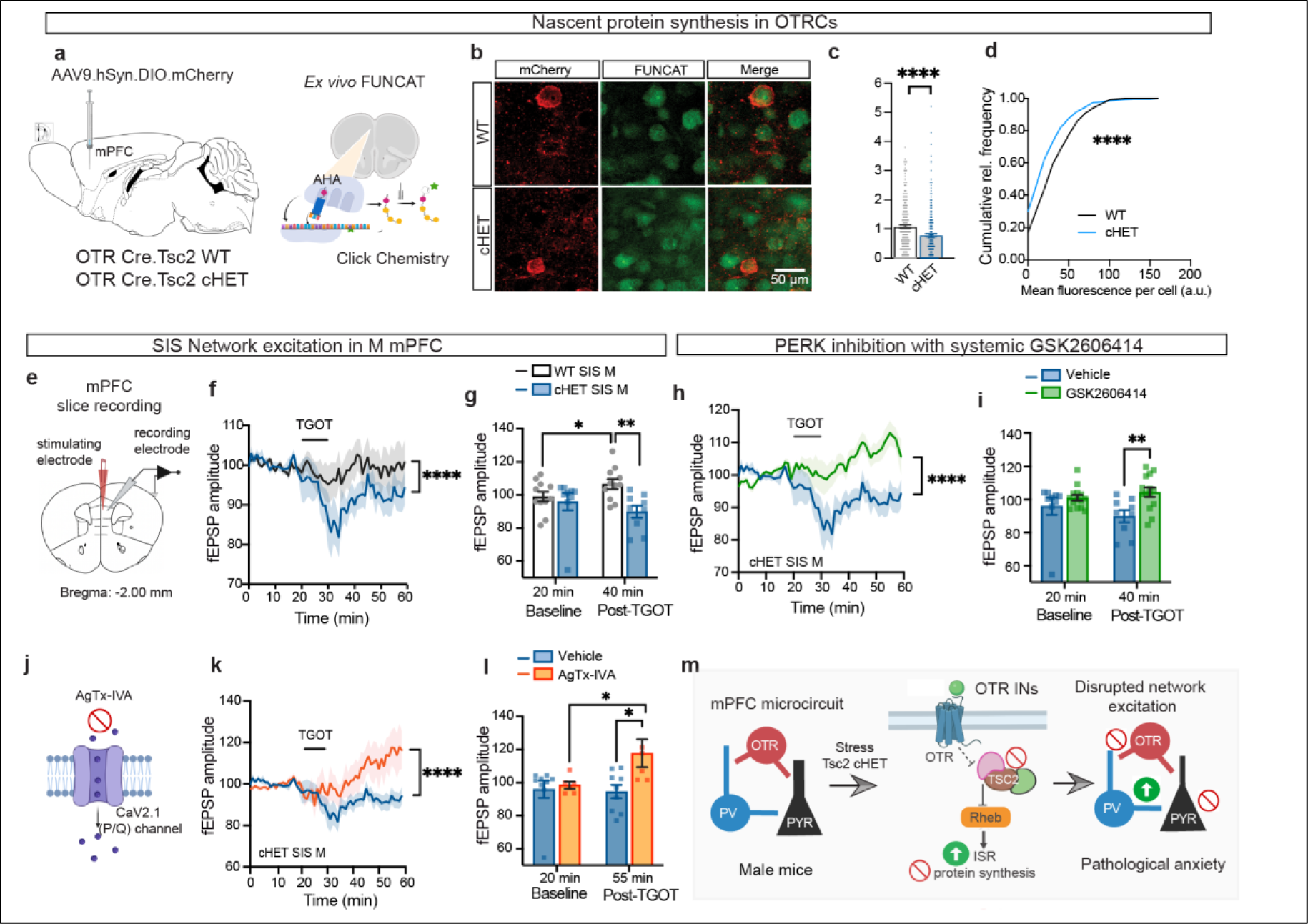
Dysfunctional OTRCs in mPFC mediate network suppression. **a)** Schematic showing viral expression of mCherry in mPFC OTRCs and the *ex vivo* fluorescent non-canonical amino acid tagging in mPFC slices of WT and cHET brains. **b)** Representative image from immunohistochemistry carried out with anti-mCherry and click chemistry-based FUNCAT label. **c)** Optical density of FUNCAT signal in OTRCs shows decreased nascent protein synthesis in cHET mice compared to WTs. **d)** Cumulative relative frequency of FUNCAT fluorescence. **e)** Schematic showing electrophysiological recording of evoked field potentials from the inner layer of mPFC (L5) while stimulating L2/3. **f)** TGOT superfusion in male brain slices leads to a sharp decrease in output spiking activity in pyramidal layer of cHET mPFC while it leads to a delayed excitation in WT mPFC. **g)** Bar plots showing TGOT-elicited increase in network excitation in WT SIS mPFC and the decline in network excitation in cHET SIS mPFC at 40 min post TGOT treatment. **h)** Subchronic PERK inhibition with GSK2606414 rescues the TGOT-elicited network suppression in male cHET mPFC. **i)** Bar graphs showing rescue of network activity at 40 min post-TGOT application. **j)** Schematic showing the action of ω-Agatoxin IVA (AgTx-IVA) in blocking P/Q-type Calcium channel. **k)** Pre-incubation of cHET mPFC slices with AgTx-IVA rescues the OT-elicited network suppression and restores delayed excitation in the inner pyramidal layer. **l)** Bar plots showing increase in network activity with TGOT application in AgTx-IVA pre-treated slices. **m)** Model of cortical network disruption in the conditional OTRC:Tsc2 cHET mouse model. Statistical tests: c) Unpaired t-test, f-I, k-l) RM Two-way ANOVA with Bonferroni post-hoc test. Sample size: c, d) n= 259 WT cells and 274 cHET cells from 4 mice/ group. f-g) n= 6-14 slices from 3-5 mice/group.

Considering activation of the ISR observed in the mPFC of cHET males, we next measured field potentials from socially isolated male mice that underwent subchronic oral administration of GSK2606414. As predicted, PERK inhibition rescued the output spiking activity of mPFC in male cHETs (**Fig. 5h, i**), but did not alter the prefrontal network activity of male WT (**Extended Data Fig. 9d, e**). These findings indicate that the network aberration caused by *Tsc2* deletion in OTRCs is mediated by the PERK-dependent ISR. The sharp decline in excitatory output from mPFC with translationally silent OTRCs in male mPFC may seem incongruent with the inhibitory role of OTR interneurons. However, it is known that SST interneurons not only form inhibitory contacts onto excitatory pyramidal neurons (PNs) in the mPFC but they also provide potent inhibitory input onto parvalbumin (PV) interneurons causing a net disinhibition of pyramidal neurons^63,64^. Thus, we assessed whether the deficient network activity is caused by the lack of OTRC mediated inhibition of PV interneurons. Given the P/Q-type calcium channel dependence of synaptic output from PV interneurons^65,66^, we pre-treated the mPFC slices with selective P/Q type Ca^2+^ channel blocker, ω-Agatoxin IVA (AgTx-IVA)^67^ (**Fig. 5j**). This strategy effectively rescued TGOT-induced network suppression in stress-exposed cHET mPFC and additionally, led to a greater network excitation indicating that major suppression of the mPFC network is mediated by PV INs (**Fig. 5k, l**). These findings support a cortical network model where OTRCs disinhibit L5 pyramidal cells by sending strong inhibitory input to the PV Ins (**Fig. 5m**). Disruption of OTRC function caused by *Tsc2* disruption is further aggravated when oxytocin signaling is recruited, either with pharmacological treatment of TGOT or by anxiogenic environments, in a high-stress state of social isolation. Because oxytocin signaling is thought to block the TSC complex, this function has an amplified effect in *Tsc2* haploinsufficient cells and leads to further increase in ISR. Therefore, at the network level, the evoked synaptic potential in output layers of mPFC is disrupted by OTRC-specific *Tsc2* depletion.

## Discussion

In this study, we investigated the interaction of genetic background rendered by *Tsc2* haploinsufficiency, and prolonged social isolation stress in anxiogenic environments. We found evidence of prefrontal network suppression caused by social isolation in mutant males, which likely contributes to the excessive avoidance and seeking of social buffering of stress (**Extended Data Fig. 10**). Our findings are consistent with prior studies showing the effects of chronic social isolation in reducing the excitability of pyramidal cells in deep layers of mPFC. Likewise, Park et al^68^ showed that sustained anxiety state evokes hypofrontality by reducing the firing rate of spontaneously active neuron subpopulations in PFC, leading to a decline in prefrontal control of flexible decision-making. Interestingly, despite activation of the same molecular pathway (ISR) in response to SIS in both male and female mPFC, only male mutants exhibit anxiety-like behaviors and mPFC suppression, whereas female mutants develop stress resilience. In this context, Li et al^31^ reported that OTR interneurons in mPFC specifically express of corticotropin-releasing-hormone-binding protein (CRHBP), which serves to antagonize the function of stress hormone CRH. Although CRHBP is expressed in both male and female OTRCs, the effect of CRH in evoking spikes in L2/3 pyramidal cells that express the receptor for CRH (CRHR1) was far greater in males than in females. In future studies, it will be interesting to explore the connection between CRHBP and hypofrontality – it is possible that CRHBP expression was downregulated during sustained ISR in OTR interneurons in male mPFC which impaired the modulation of L2/3->L5 synaptic transmission, resulting in an unchecked stress response.

Our findings indicate that cell type-specific *Tsc2* heterozygous deletion leads to acceleration of UPR response via PERK arm, which then precipitates maladaptive avoidance and loss of motivational drive in male mice in response to subthreshold social isolation stress for 10 days. The connection between chronic social isolation and ISR was previously reported in a study which showed that chronic social isolation in fruitflies causes fragmented sleep and elicit unfolded protein response (UPR) by activating all three major pathways of UPR, including increase in BiP, splicing of Xbp1, and phosphorylation of eIF2α (Ser51) via PERK activation^69^. Of note, hyperactivation of mTORC1 can also lead to increase in endoplasmic reticulum (ER) stress via uncontrolled protein synthesis and proteotoxic load. Although we observed an increase in mTORC1 activation in OTRCs, this effect was neutralized by PERK-ISR pathway, thereby causing a net decline in protein synthesis. Thus, it is unlikely that the ISR in OTRCs was due to mTORC1-mediated proteotoxicity. In addition, rapamycin was only able to rescue the motivational drive in mutant males and did not correct anxiety-related avoidance of the center zone in open field indicating that the mTORC1 and PERK arms are activated in parallel. Overall, our findings demonstrate that cell-type specific Tsc2 haploinsufficiency in OTRCs modifies the susceptibility to social isolation. Given the high prevalence of anxiety disorders, it is important to investigate the subtypes of pathological anxiety that have distinct circuit-based molecular mechanisms.

## Methods

### Animals

Mice were provided with food and water *ad libitum* and were maintained in a 12h/12h light/dark cycle at New York University or Stony Brook University at stable temperature (78°F) and humidity (40 to 50%). All mice were backcrossed to C57Bl/6J strain for at least 5 generations. Both male and female mice, aged 3-6 months, were used in all experiments. Floxed Tsc2 mice were kindly provided by Dr. Michael Gambello (Emory University). OTR Cre BAC transgenic mice (Founder line #ON66) were generated by GENSAT and kindly provided by Dr. Nathaniel Heintz (The Rockefeller University). Wildtype C57Bl/6J mice (stock #000664) were purchased from Jackson labs. All procedures involving the use of animals were performed in accordance with the guidelines of the National Institutes of Health and were approved by the University Animal Welfare Committee of New York University and Stony Brook University.

### Drugs and chemicals

Rapamycin stock (50 mg/mL) was dissolved to 1 mg/mL in 5% Tween-80, 15% polyethene glycol 400 (PEG-400), and 0.9% saline. Rapamycin at 5mg/kg (LC Laboratories #R-5000) was injected intraperitoneally for 3 days. PERK inhibitor GSK2606414 (MedChem Express #HY-18072) was first dissolved in DMSO to prepare the stock concentration of 100 mg/ml. The GSK2606414 stock was diluted to 10 mg/ml with 0.5% HPMC, 0.1% Tween-80 and ddH_2_O, and administered to mice at 50 mg/kg by oral gavage, once a day for 3 days. Thr^4^, Gly^7^-Oxytocin (TGOT) (Bachem, # 4013837) was dissolved in ddH_2_O at a concentration of 0.4 mM for stock solution and was further diluted in sterile saline to a final concentration of 16 μM for slice electrophysiology. ω-Agatoxin IVA was dissolved in ddH_2_O to prepare 5 µM stock, which was further delivered in the bath for a final concentration of 0.5 µM. Azidohomoalanine (AHA) (Click Chemistry Tools, #1066-100) was dissolved in ddH_2_O at a stock concentration of 100 mM and diluted to 1 mM in sterile saline. Stock solution of aqueous 32% paraformaldehyde (EMS, #15714) was freshly diluted to 4% in 0.1 M PBS for transcardial perfusions and post-fixation of brain slices.

### Stereotaxic surgeries

Mice were anesthetized with the mixture of Ketamine (100 mg/kg) and Xylazine (10 mg/kg) in sterile saline (i.p. injection). Adequate sedation was determined by a lack of gentle toe pinch withdrawal reflex. A lubricant eye ointment (Genteal Tears; Alcon) was applied on the eyes to prevent ocular dehydration. Visual monitoring of respiration occurred periodically throughout surgery to confirm survival. Stereotaxic surgeries were carried out inside a Class II A2 biosafety cabinet (LabRepCo, # NU-677-500) on the Kopf stereotaxic instrument (Model #942), which was equipped with a Nanojector UMP3TA (WdI). Viral vectors were injected intracranially using 2.0 μl Neuros syringe (Hamilton, #65459-02) at 1 nl/s. Before removing the needle, an interval of 10 min was allotted for viral diffusion. Upon completion of intracranial injections, the incision of the scalp was closed using a tissue adhesive (3M Vetbond). Normothermia during surgery was maintained at 36.6°C by resting the mouse on a covered heating pad (7 cm X 7 cm) connected to a rodent warmer console (Stoelting, # 53800M). To prevent somatic dehydration, each mouse was subcutaneously injected with 500 μl of sterile saline before surgery, and further provided with hydrogel (Clear H2O, # 70-01-5022) in the recovery cage. General monitoring and analgesia using 3 mg/kg subcutaneous Ketoprofen (Zoetis) were provided up to three days post-surgery. To knock down Rheb in OTRCs, 200 nl of lenti pSICO.DIO.shRheb (1.00 X 10^13^ GC/ml, packaged by Vigene, Addgene #81087) was bilaterally injected into the medial prefrontal cortex [-2.00 mm anterior posterior (AP), +/-0.25 mm mediolateral (ML) and -1.60 mm dorsoventral (DV)] of either OTR Cre^+/-^ wildtype od OTR Cre^+/-^.Tsc2^f/+^ (cHET) mice. The same mice also went through a prior surgery three weeks earlier with AAV9.hSyn.DIO.mCherry.WPRE (1.00 X 10^13^ GC/ml; Addgene #50459) to label OTRCs in mPFC, since the pSICO vector has no fluorophore expressed in Cre+ cells that are targeted for Rheb knockdown.

### Behavior

All behavior sessions were conducted during the light cycle (7:00 AM – 7:00 PM EST). Both male and female mice were included in behavior experiments. Tsc2 cHET and control mice were randomly assigned to experimental conditions including drug and vehicle administration, and for the order of testing in any given paradigm. Behavior experiment data were collected by experimenters that were blind to the experimental conditions and genotype. Following weaning, mice were group-housed with up to three sex-matched littermates in standard cages (8 in X 13 in X 5 in) fitted with corn cob bedding and Enviro-dri enrichment. Acclimation of the mice in the behavior room lasted at least 30 min prior to testing. A white noise machine (Yogasleep Dohm) was used to blunt any environmental noise that may induce confounding anxiety. All items in the behavior equipment were cleaned with 30% ethanol unless otherwise stated. Behavior testing occurred in the order of least to most demanding to the mouse subjects.

### Social Isolation stress

Adult mice (males and females, ∼3 months old) were removed from their group-housed cage and singly housed in a new standard cage for the remainder of their life-span. They were provided with food and water *ad libitum* and maintained in a 12h/12h light dark cycle at a stable temperature (78^0^F) and humidity (40 – 50 %). The cages were provided with Enviro-dri enrichment identical to group-housed mice. Following 10 days of social isolation stress, mice were subjected to a series of behavioral experiments or euthanized to perform histology and slice electrophysiology.

### Feeding assay

Singly housed mice were food-deprived for 24h, following which they were provided with a single chow pellet in the center of their home cage. The amount of chow consumed by the animal over 10 min was manually weighed and recorded by an experimenter blind to the genotype.

### Open field activity

Mice were placed in the center of an open field (27.31 cm x 27.31 cm x 20.32 cm) for 15 min during which a computer-operated optical system (Activity monitor software, Med Associates) monitored the spontaneous movement of the mice as they explored the arena. The arena was divided into center zone (inner square: 13.67 cm X 13.67 cm) and total zone encompassing the whole arena of open field. The parameters measured were distance traveled, and the ratio of distance traveled in the center to that in the total zone. Further, the data was analyzed as distance traveled across three time-bins, 5 min each.

### Elevated Plus Maze

Mice were allowed to explore the arms of a plus maze elevated 38.5 cm above the floor for 5 min. The open arms and closed arms were each 5 cm X 30.5 cm whereas the center zone was 5 cm X 5 cm. A video tracking software (Ethovision XT15) was used to measure the duration and frequency of entries into the open and closed arms.

### Marble burying test

Mice were placed into a standard polycarbonate rat cage (28 cm x 19 cm x 15 cm) equipped with 2 inches of fresh bedding (ALPHA-dri) and 20 black glass-marbles (14 mm diameter) arranged as a matrix with 4 rows and 5 columns. The video of the trials was captured using a computer-operated video-tracking software (Ethovision XT15). Each trial was carried out for 30 min. The number of marbles buried every 5 min and the latency to bury the first marble were manually scored.

### Self-grooming test

Self-grooming test was conducted in 18 cm X 18 cm X 30 cm cuboid chambers (Coulbourn instruments) housed inside sound-attenuated cubicles. Lighting in the chamber was provided with white houselight, and the floor of the chamber was covered with a white Plexiglas platform. No enrichment was provided, to suppress exploration and to encourage grooming behavior. Each trial was conducted for 30 min. A video-recording of grooming behavior was captured using USB video cameras (AM2-CA01, Actimetrics) and manually scored for latency to groom and total grooming time in the last 10 min of the session.

### Three-Chamber Social Interaction Test

Each mouse was introduced into a 3-chamber social box (Harvard Apparatus). The arena was divided into three chambers – two 42.5 x 19.5 x 23 cm side chambers (left and right) and one 42.5 x 18 x 23 cm center chamber with two 10 (W) x 23 (H) cm removable doors that separate the center chamber from the side chambers. Each side chamber had a cylindrical wire enclosure placed in the outer corner. In the first trial, the enclosures were empty, and the subject mouse was allowed to freely explore the arena spanning three chambers. In the second trial, an enclosure in one of the lateral chambers had a novel object whereas the other enclosure in the contralateral chamber had a sex-matched stranger mouse. The locations of the novel object and mouse were counterbalanced across trials. The subject mouse was allowed to freely explore the social and non-social targets. Each trial ran for 5 min. A video-tracking software was used to track the movements of the mice (Ethovision XT15). The duration and frequency of the subject mouse in the zone of interaction (1 inch radius outside of the footprint of enclosure) were assessed. Sociability index was calculated as follows:

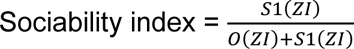

Where S1(ZI) is the duration of the subject mouse in stranger mice zone of interaction, and O(ZI) is the duration of the subject mouse in the object zone of interaction.

### Resident-Intruder Test

For the resident-intruder test, an unfamiliar male BALB/cJ mouse of similar body weight and age was introduced into the home cage of the resident animal for 10 min. The enrichment, as well as food and water bottle were removed from the cage during the experiment. The dyadic interaction between the resident mouse and intruder in the rearing cage were video-recorded using GigE USB camera and Ethovision XT13 software. Following parameters were manually scored - the amount of time that the resident animal spends investigating the intruder (following, sniffing, grooming) and the amount of time spent on physical attacks (biting, clawing, wrestling, and chasing).

### Mating Test

For the mating test, a sexually receptive C57Bl/6J female in estrus phase was selected as the mating partner and introduced into the home cage of the male subject for 10 min. The enrichment, as well as food and water bottle were removed from the cage during the experiment. The dyadic interaction between the mice were video-recorded using GigE USB camera and Ethovision XT13 software. An experimenter blind to the genotype manually scored the time the male subject spends investigating the female mouse (following, sniffing, grooming) and the amount of time spent on mounting and consummatory behaviors.

### Slice Electrophysiology

Fresh brain slices from conditional heterozygote mice (OTR Cre^+/-^.Tsc2^f/+^cHET) and their wildtype littermates were used for ex vivo slice electrophysiology experiments. All mice were singly housed for at least 10 days to induce social isolation stress before the recordings. Mice were deeply anesthetized with halothane and perfused with ice-cold slicing solution (85 mM NaCl, 2.5 mM KCl, 1.2 mM NaH_2_PO_4_, 25 mM NaHCO_3_, 0.5 mM CaCl_2_, 7 mM MgSO_4_, 25 mM glucose, 75 mM sucrose, and 0.5 mM ascorbic acid). The brain was then decapitated and extracted into a beaker filled with ice-cold slicing solution equilibrated with 95% O_2_/5% CO_2_. The chilled brains were blocked at the level of the optic chiasm and sectioned coronally from rostral to caudal into 300 μm slices using a vibratome. A selected coronal PFC slice was maintained on an interface chamber at 31°C for at least 1 h with artificial cerebrospinal fluid (ACSF) flow rate of 1.5–2.5 ml/min for recording. The ACSF contained 125 mM NaCl, 3.3 mM KCl, 1.2 mM NaH_2_PO_4_, 25 mM NaHCO_3_, 2 mM CaCl_2_, 1.2 mM MgSO_4_, 15 mM glucose, and 0.5 mM ascorbic acid. A glass recording electrode (filled with ACSF, 5 MΩ resistance) was guided to the inner layers (L5) of the medial prefrontal cortex, and a bipolar stainless steel stimulating electrode (75 kΩ) was positioned to target layer 2/3 of mPFC. Orthodromic synaptic potentials were evoked via an isolated current generator (100 μs pulses of 0.3–0.7 mA; Digitimer). Evoked field potentials were recorded with an Axoclamp 2B amplifier and Axon WCP software (Molecular Devices). Field EPSP (fEPSP) was measured as a change in evoked field potential amplitude. Baseline responses were recorded for 20 min at 0.05 Hz with a stimulus intensity of 40–50% of maximum fEPSP. When stated, slices were pre-incubated with 0.5 µM ω-Agatoxin IVA. At the 20 min mark of the baseline recording, a selective oxytocin receptor agonist, [Thr4,Gly7]-oxytocin (TGOT) (2 µM) (Bachem, #4013837), was superfused for 10 min with the same intensity and pulse duration as the baseline stimuli. fEPSP was recorded continuously for 30 min thereafter. Data were analyzed offline using WCP PeakFit (Molecular Devices). The n refers to the slice number. All fEPSP data are presented as mean ± standard error of the mean (SEM). Average traces were calculated over 1 min (n = 6, consecutive sweeps) of stimulation and plotted. These averaged traces were normalized to the mean peak amplitude that was recorded over the 20 min baseline. Experimental groups were compared using a two-tailed t-test.

### Fluorescent non-canonical amino acid tagging (FUNCAT)

300 μm-thick brain slices containing medial prefrontal cortex (mPFC) [Bregma +2.34 mm to +1.94 mm] were prepared in cold (4°C) carboxygenated (95% O_2_, 5% CO_2_) cutting solution (110 mM sucrose, 60 mM NaCl, 3 mM KCl, 1.25 mM NaH_2_PO_4_, 28 mM NaHCO_3_, 5 mM Glucose, 0.6 mM Ascorbate, 7 mM MgCl_2_ and 0.5 mM CaCl_2_) using a VT1200S vibratome (Leica). Slices were recovered in oxygenated ACSF solution (125 mM NaCl, 2.5 mM KCl, 1.25 mM NaH_2_PO_4_, 25 mM NaHCO_3_, 25 mM glucose, 1 mM MgCl_2_ and 2 mM CaCl_2_) at 32°C for 2 h. 1 mM AHA was then added to the ACSF solution for an additional 2.5 h to label nascent protein synthesis. Following incubation, the mPFC was dissected out and submerged in 4% paraformaldehyde (EMS) at 4°C overnight. The fixed micro-slices were embedded in 3% agarose, re-cut into 40 μm sections using a vibratome, and stored at -20 °C in the cryosolution overnight. The mPFC sections were blocked and permeabilized in 5% IgG protease-free BSA, 3% normal goat serum, 0.3% Triton X-100 in 1X PBS at RT for 90 min with agitation. The brain sections were then subjected to click chemistry using Cell Reaction kit (Thermo Fisher, #C10269) with 25 μM Alkyne Alexa fluor 405 (Click Chemistry tools, #1309-1), CuSO_4_ and additive overnight at 4°C. The following day, the sections were washed 3 times with 1X PBS for 10 min per wash. Sections were blocked with 1% normal goat serum (NGS) in 1X PBS for 1 h followed by incubation with primary antibodies [rat anti-mCherry (Thermo Fisher, #M11217)] overnight at 4°C for standard immunohistochemistry.

### Immunohistochemistry

Mice were deeply anesthetized with a mixture of ketamine (150 mg/kg) and xylazine (15 mg/kg), and transcardially perfused with 0.1M PBS followed by 4% paraformaldehyde (EMS) in PBS. Brains were removed and postfixed in 4% PFA for 24h. The PFA is then replaced with 30% sucrose solution for another 24 h. 40 μm free-floating coronal brain sections containing PFC were collected using Leica vibratome (VT1000s) and stored in 1X PBS containing 0.05% sodium azide at 4°C. After blocking in 5% normal goat serum in 0.1 M PBS with 0.1% Triton X-100, brain sections were probed overnight with primary antibodies [chicken anti-EGFP (Abcam #ab13970), rabbit anti-EGFP (Thermo Fisher #G10362), rabbit anti-p-eIF2α S51 (Cell Signaling #9721), mouse anti-NeuN (Millipore Sigma #MAB377), rat anti-mCherry (Thermo Fisher #M11217), rabbit anti-p-S6 (240/244) (Cell Signaling #5364S), mouse anti-p-S6 (S235/6) (Cell Signaling # 62016S), rabbit anti-RHEB (Fisher #PIPA520129) and rabbit anti-TSC2 (Sigma #HPA030409). After washing three times in 0.1M PBS, brain sections were incubated with Alexa Fluor conjugated secondary antibodies 1:200 (Thermo Fisher #A-11001, #A32931, #A32733, #A48255, #A48254, #A32740) in blocking buffer for 1.5h at RT and mounted using Prolong Gold antifade mountant with or without DAPI (Fisher # P36931, # P36930).

### Image acquisition and analysis

Imaging data from immunohistochemistry experiments were acquired using a laser scanning microscopy (LSM) confocal microscope (Zeiss) with 20X objective lens (with 1X or 2X zoom) and z-stacks (approximately 6 optical sections with 0.563 μm step size) for three coronal sections per mouse from AP Bregma +1.98 mm to + 1.78 mm (n = 3 mice) were collected. Imaging data was analyzed with ImageJ using the Bio-Formats importer plugin. Maximum projection of the z-stacks was generated followed by manual outline of individual cells and mean fluorescence intensity measurements using the drawing and measure tools. Mean fluorescence intensity values for all cell measurements were normalized to the mean fluorescence intensity for controls. In the case of verifying Rapamycin’s effect on mTORC1 pathway in mPFC, field fluorescence of the entire image area was measured, instead of outlining each cell. This was done to prevent underrepresentation of images with fewer labeled cells. Each image was opened with ImageJ, and the 6 z-stacks of the images were subjected to maximum projection with maximum intensity. A rectangle was used to outline the entire perimeter of the image, and the fluorescence was measured. The area, and mean intensity of the image were measured.

### Statistics

Statistical analyses were performed using GraphPad Prism 8 (GraphPad software). Data are expressed as mean +/-SEM. Data from two groups were compared using two-tailed unpaired Student’s t test. Multiple group comparisons were conducted using one-way ANOVA, two-way ANOVA, or RM Two-way ANOVA with post hoc tests as described in the appropriate figure legend. Statistical analysis was performed with an α level of 0.05. P values <0.05 were considered significant.

## Acknowledgements

We thank A. Nnenna Chime, N. Pena, Chien-Yu Juan, M. Singh, and S. Chu for technical assistance; M. Nakajima and Dr. N. Heintz for the OTR Cre ON66 BAC transgenic mice; Dr. M. Gambello for the floxed Tsc2 mouse strain and A. Bordey for the pSICO.shRheb lentiviral plasmid. We are grateful to all members of the Klann and Shrestha laboratories for feedback and discussions. This study was supported by National Institute of Health grants NS034007 and NS047384 to E.K., and MH132795 to P.S. P.S. is also supported by a Sloan research fellowship 94485.

## Extended Data Figures

**Extended Data Figure 1.**
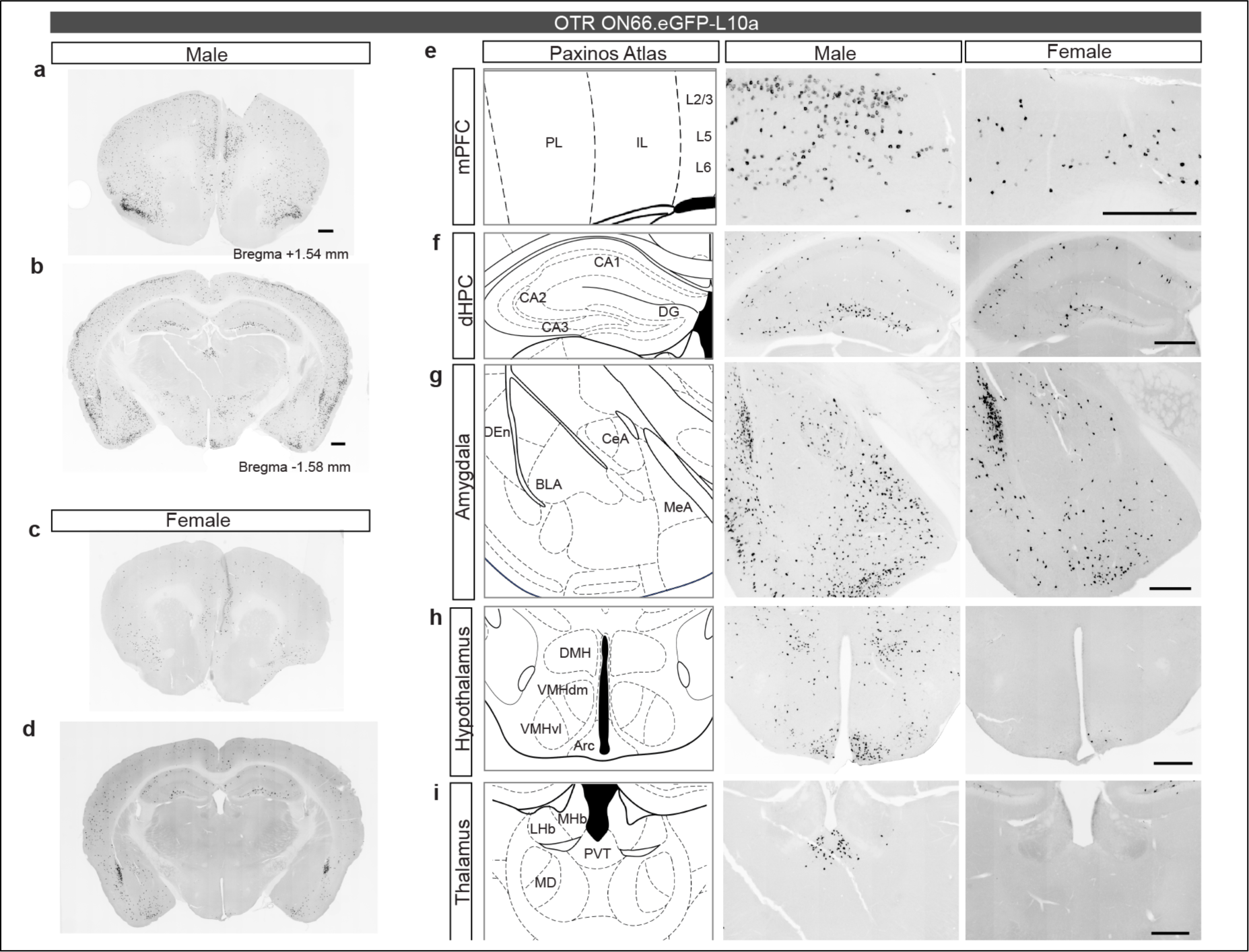
Cre recombinase activity in male and female brains for double transgenic OTR ON66.eGFP-L10a mouse strain. **a)** Gene expression pattern for GFP-tagged ribosomal protein (GFP-L10a) in the anterior brain section (Bregma +1.54 mm) containing mPFC of male brain **b)** Gene expression pattern for GFP-L10a in the posterior brain section (Bregma -1.58 mm) containing dorsal hippocampus, paraventricular thalamus, hypothalamus and amygdala of male brain. **c)** Gene expression pattern for GFP-L10a in the anterior brain section containing mPFC of female brain. **d)** Gene expression pattern for GFP-L10a in the posterior brain section of female brain. **e)** Reporter gene expression in the male mPFC encompasses the superficial layer (L2/3) and inner layer (L5) whereas the female mPFC has cells labeled only in inner layer (L5). **f)** Reporter gene expression in dorsal hippocampus (dHPC) shows graded pattern with higher expression in dentate gyrus and low expression in CA1 in males, and uniform expression across the subregions of dHPC in females. **g)** GFP-L10a expression is greater in the medial amygdala of males than in females, but otherwise the pattern of gene expression is similar in amygdala for basolateral amygdala and central amygdala. **h)** Reporter gene expression in the male hypothalamus shows robust enrichment in the Arcuate nucleus (Arc) and sparse expression in ventromedial hypothalamus (VMHvl). GFP-L10a is sparsely expressed only in Arc nucleus of female hypothalamus. **i)** GFP-L10a is expressed strongly in the paraventricular nucleus (PVN) whereas this is absent in the female PVN. Scale bar = 50 μm.

**Extended Data Figure 2.**
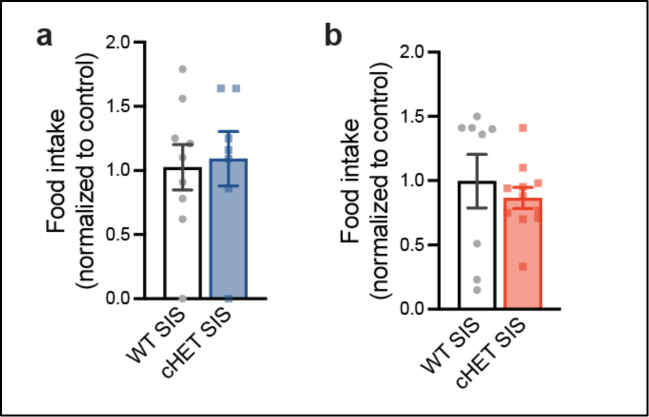
Feeding behavior in conditional TSC mouse model. **a)** Food intake is comparable in both socially isolated WT and cHET male mice. **b**) Food intake is also normal in socially isolated WT and cHET females. Statistical test: Unpaired t-test. Sample size: n = 8-11 mice/group.

**Extended Data Figure 3.**
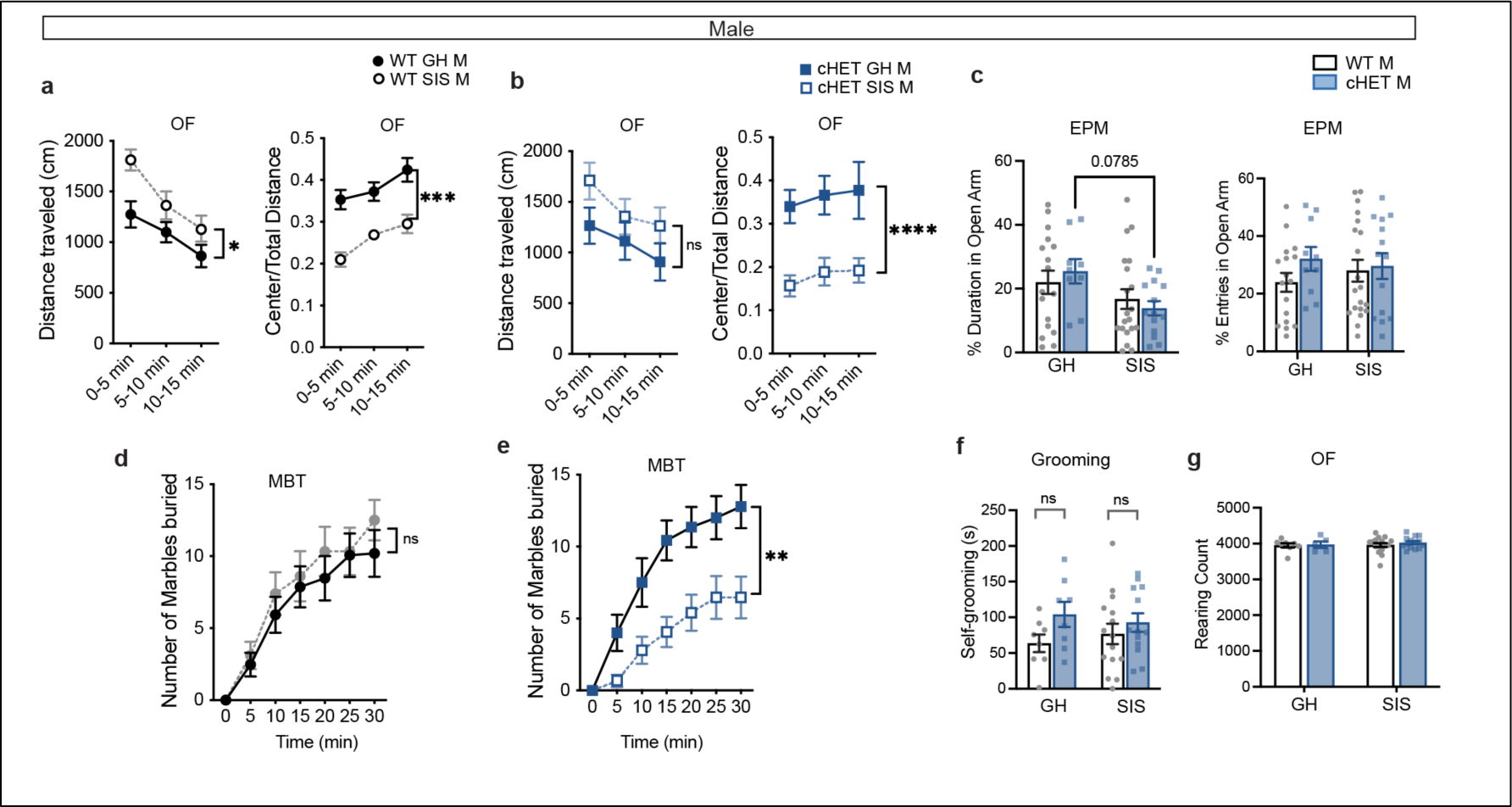
Effects of social isolation stress in male emotional and stereotypical behaviors. **a)** In WT males, SIS leads to increased exploration in Open Field, and reduces center zone approach compared to grouphoused condition. **b)** SIS does not cause an increase in spontaneous locomotion of cHET males but it strongly reduces center zone approach in open field. **c)** Compared to grouphoused condition, cHET male mice have a trend-level decrease in relative time spent in the open arm of elevated plus maze, while the WT males have no change in their behavior. Percent open arm entries in the EPM is comparable for all groups of male mice. **d)** Marble burying behavior is not affected in SIS-exposed WT males. **e)** SIS decreases marble burying behavior in cHET males. **f)** Normal self-grooming behavior is observed in all male groups. **g)** Rearing behavior in Open Field is comparable for grouphoused and singly-housed mice of both genotypes. Statistical tests: a, b, d, e) RM Two-way ANOVA; c, f, g) Two-way ANOVA with Bonferroni post-hoc test. *p<0.05, **p<0.01, ***p<0.001, ****p<0.0001, ns not significant. n=8-18 mice/group.

**Extended Data Figure 4.**
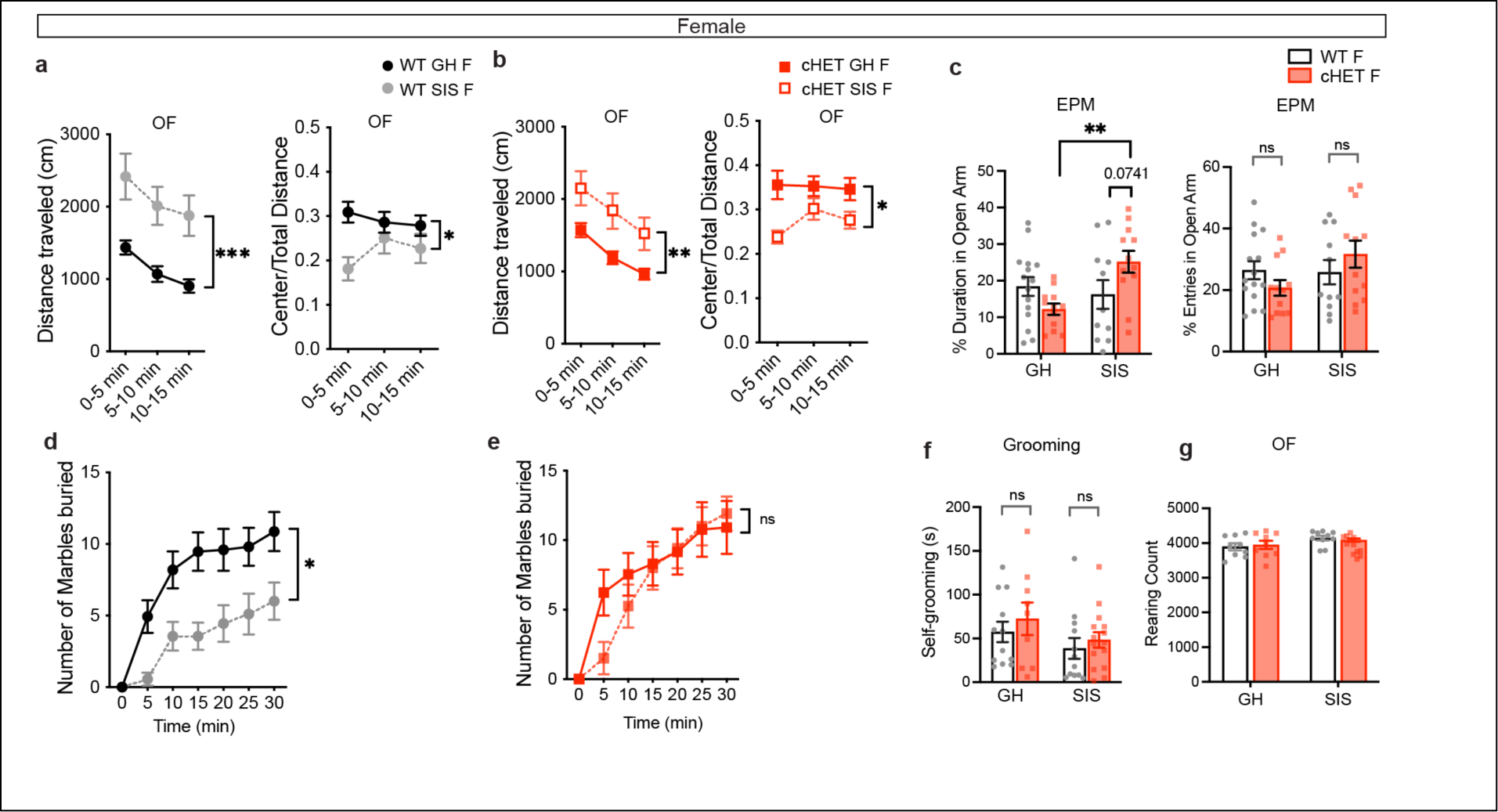
Effects of social isolation stress in female emotional and stereotypical behaviors. **a)** In WT females, SIS leads to a robust increase in spontaneous locomotion in Open Field and also reduces center zone approach in the initial time bin (0-5 min). **b)** SIS similarly increases exploratory behavior in OF and also increases thigmotaxis in cHET females in the initial time bin (0-5 min) which gradually returns to normal levels over time. **c)** cHET female mice spend significantly greater time in the open arm of EPM compared to grouphoused condition and they also have a trendlevel increase in open arm duration compared to the singly-housed WT females. Percent open arm entries is comparable in all groups of female mice. **d)** SIS-exposed WT females have reduced motivational drive to bury marbles. **e)** SIS has no effect on marble burying behavior of cHET female mice. **f)** Self-grooming behavior is normal in both WT and cHET female mice in grouphoused as well as singly-housed condition. **g)** Rearing behavior is normal in all groups of female mice. Statistical tests: a, b, d, e) RM Two-way ANOVA; c, f, g) Two-way ANOVA with Bonferroni post-hoc test. **p<0.01, **p<0.01, ***p<0.001, ns not significant. n=8-20 mice/group.

**Extended Data Figure 5.**
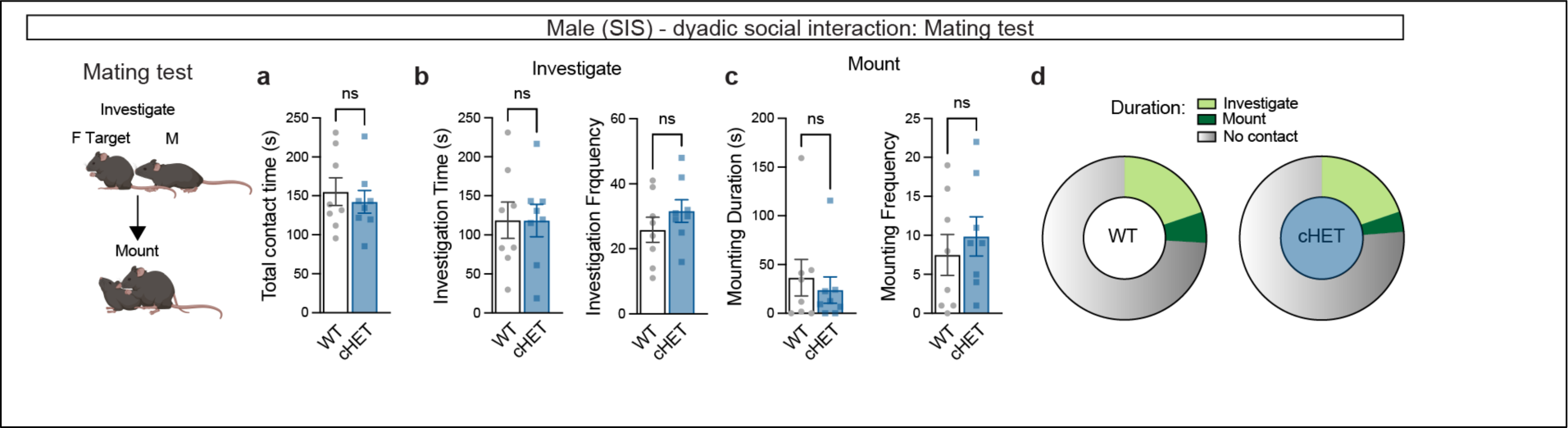
Reproductive behavior in socially-isolated male mice. **a)** Total contact time between the male subject and sexually receptive female intruder is comparable for both WT and cHET males. **b)** Investigation time (sniffing and allogrooming) and investigation frequency are normal for both WT and cHET SIS males. **c)** Mounting duration and mounting frequency are similar for both WT and cHET SIS males. **d)** Venn Diagram showing relative time spent investigating and mounting the mating partner for WT and cHET males. Statistical tests: a-c) Mann Whitney U test, ns not significant. n = 8 mice/ group.

**Extended Data Figure 6.**
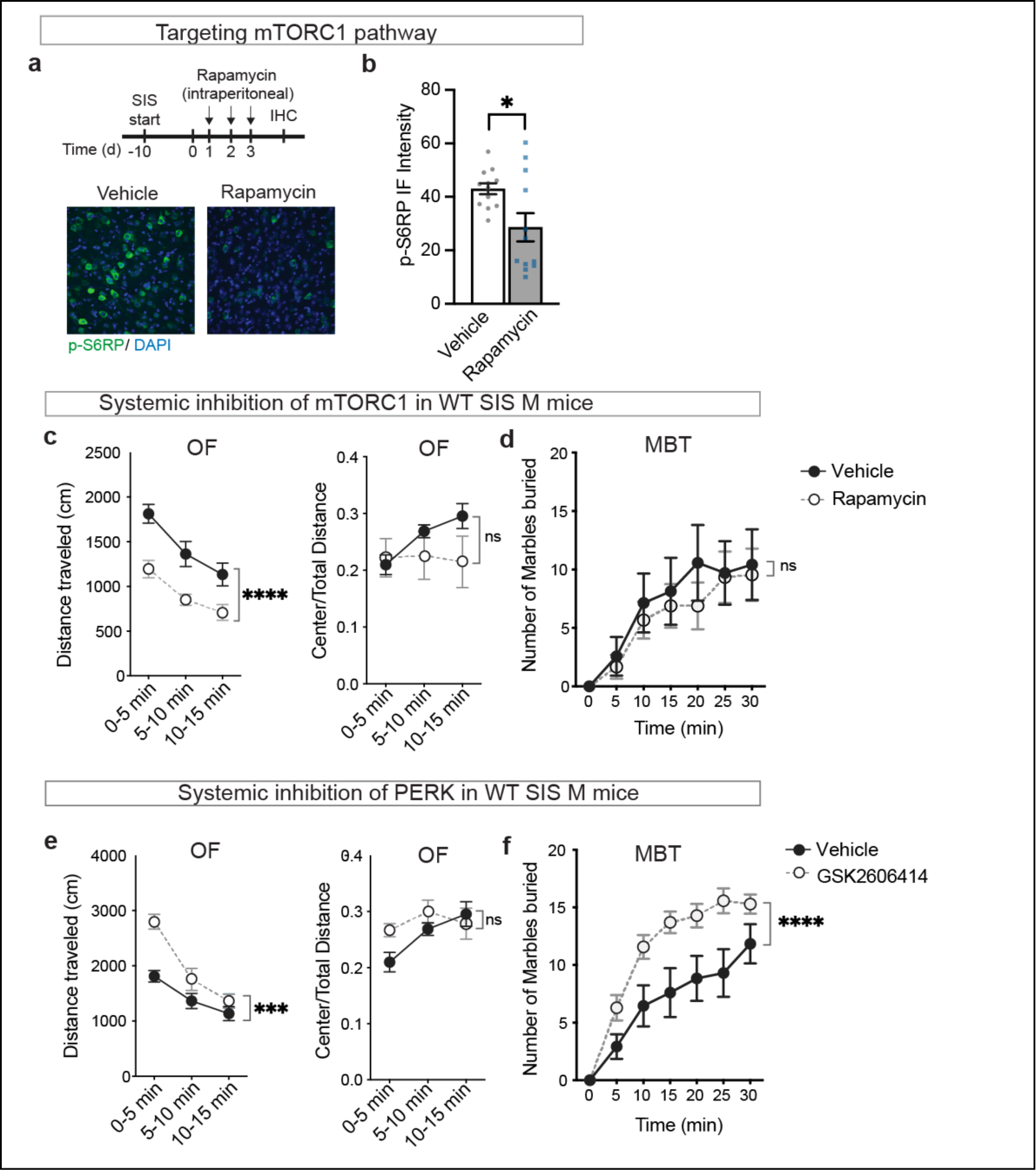
Systemic inhibition of TSC downstream effectors. **a)** Schematic for subchronic injection of Rapamycin for 3 days. Representative images for mPFC sections immunostained with anti-pS6RP with DAPI nuclear counterstain. **b)** Optical density of pS6RP immunofluorescence is significantly reduced in Rapamycin treated mPFC sections. **c)** Systemic injection of Rapamycin reduced hyperactivity in SIS-exposed WT males but did not alter thigmotaxis. **d)** Rapamycin did not alter marble burying behavior in WT SIS males. **e)** Systemic inhibition of PERK increased hyperactivity in the open field and did not affect thigmotaxis. **f)** PERK inhibition stimulated more marble burying behavior in singly-housed WT males. Statistical tests: a) Unpaired t-test; c-f) RM Two-way ANOVA with Bonferroni post-hoc test. *p<0.05, ***p<0.001, ****p<0.0001, ns not significant. n=5-17 mice/ group.

**Extended Data Figure 7.**
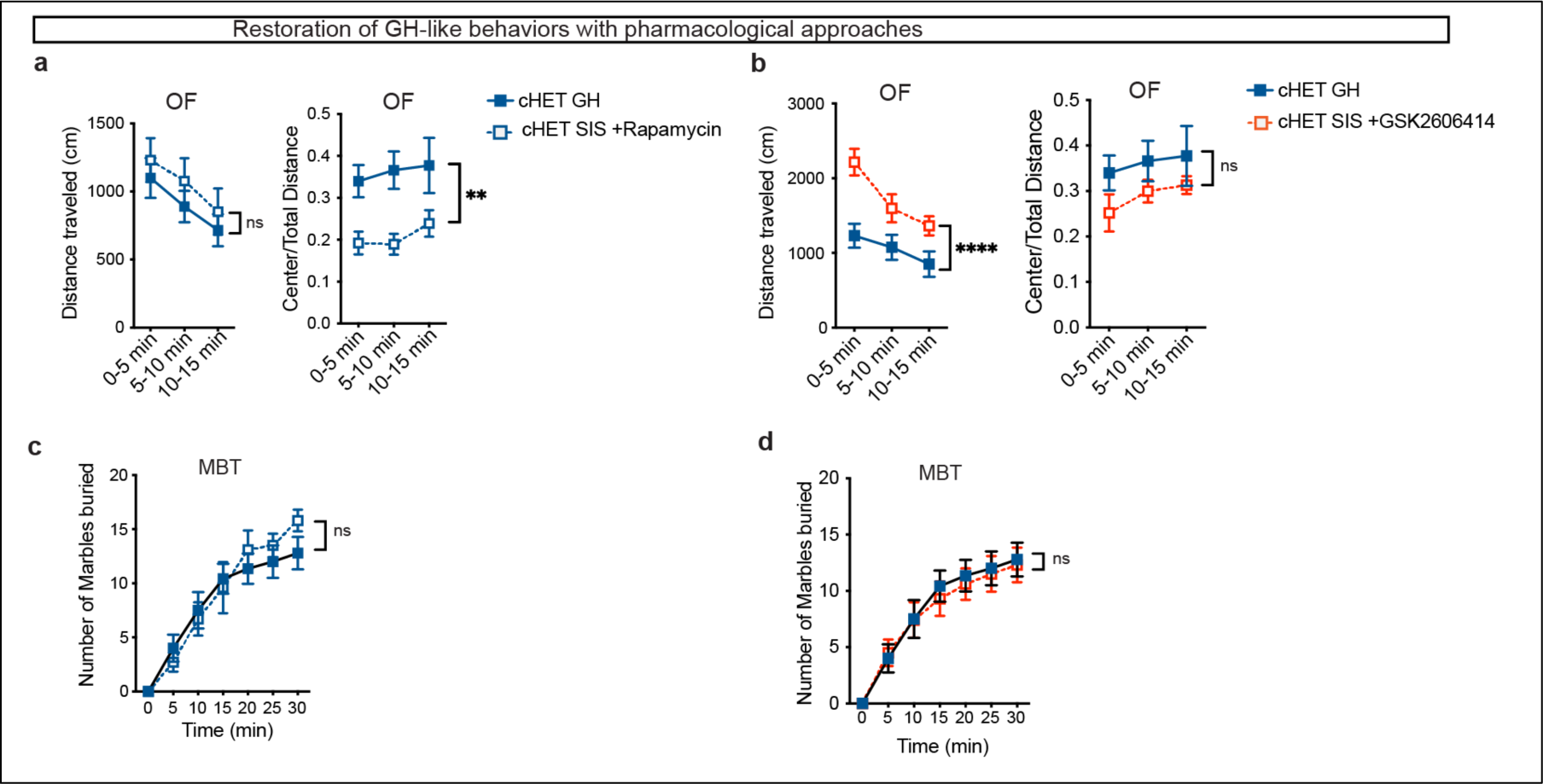
Restoration of GH-like behaviors with systemic inhibition of TSC downstream effectors. **a)** Spontaneous exploration in OF in both WT and cHET SIS-exposed males is normalized to GH-levels by Rapamycin treatment, while there is no change in thigmotaxis. **b)** PERK inhibitor GSK2606414 increases spontaneous activity in both WT and cHET males above GH-levels, but restores thigmotaxis behaviors. **c)** Marble burying behavior in cHET SIS males is restored to GH-levels with Rapamycin treatment. **d)** PERK inhibition normalizes marble burying behavior in singly-housed cHET males but it elicits stereotypy and increased marble burying in SIS-exposed WT males above GH-levels. Statistical tests: a-d) RM Two-way ANOVA with Bonferroni post-hoc test. N=6-14 mice/group.

**Extended Data Figure 8.**
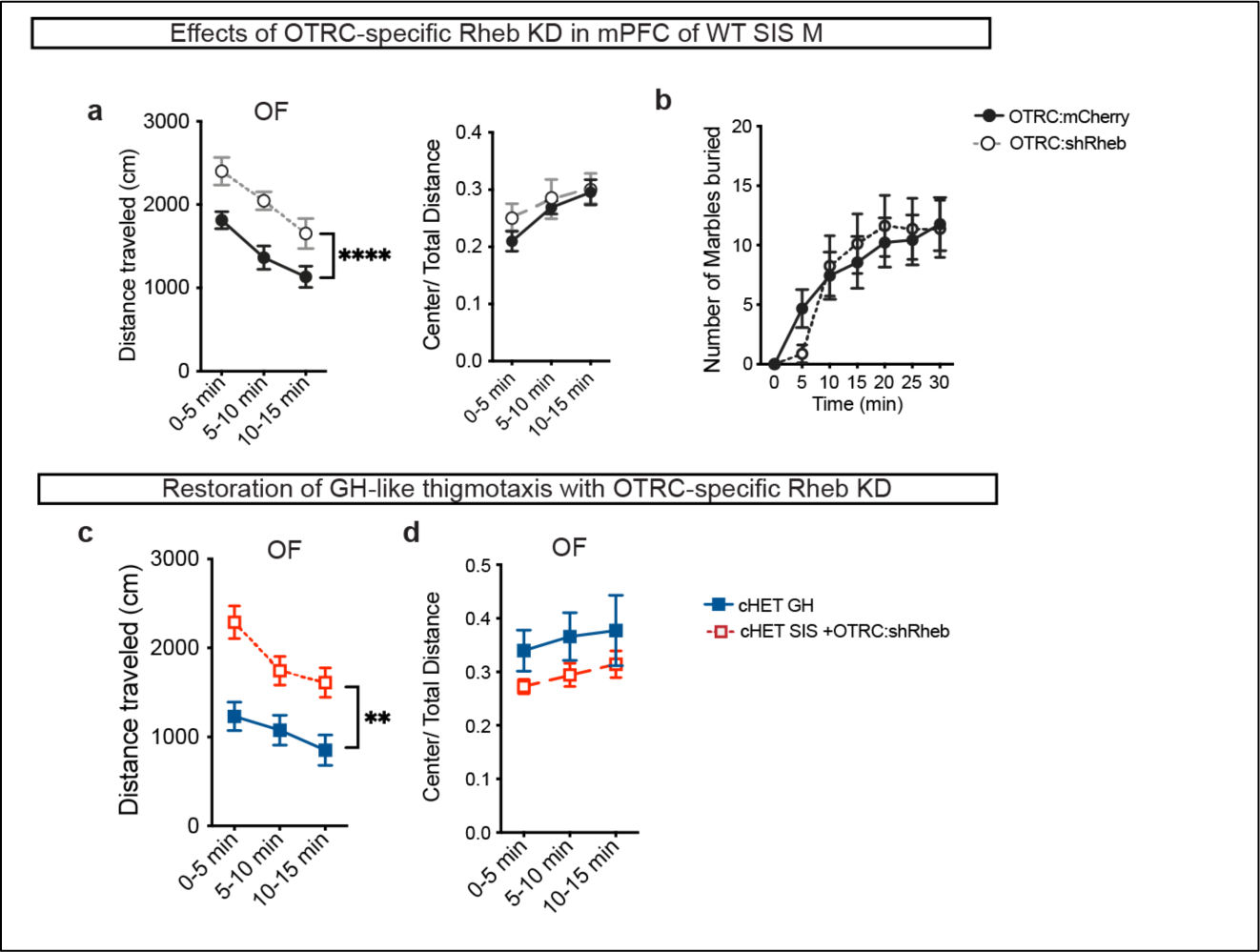
Rheb Knockdown in OTRCs in SIS-exposed WT mice. **a)** Rheb KD in prefrontal OTRCs in WT males increases spontaneous locomotion but does not alter thigmotaxis. **b)** Rheb KD does not alter marble burying behavior in SIS-exposed WT male mice. **c)** OTRC-specific Rheb KD in cHET males increases spontaneous locomotion compared to grouphoused cHET males. **d)** Rheb KD restores group-housed like thigmotaxis in socially isolated cHET male mice. Statistics: RM Two-way ANOVA.**p<0.01, ****p<0.0001. n=8-9 mice/group

**Extended Data Figure 9.**
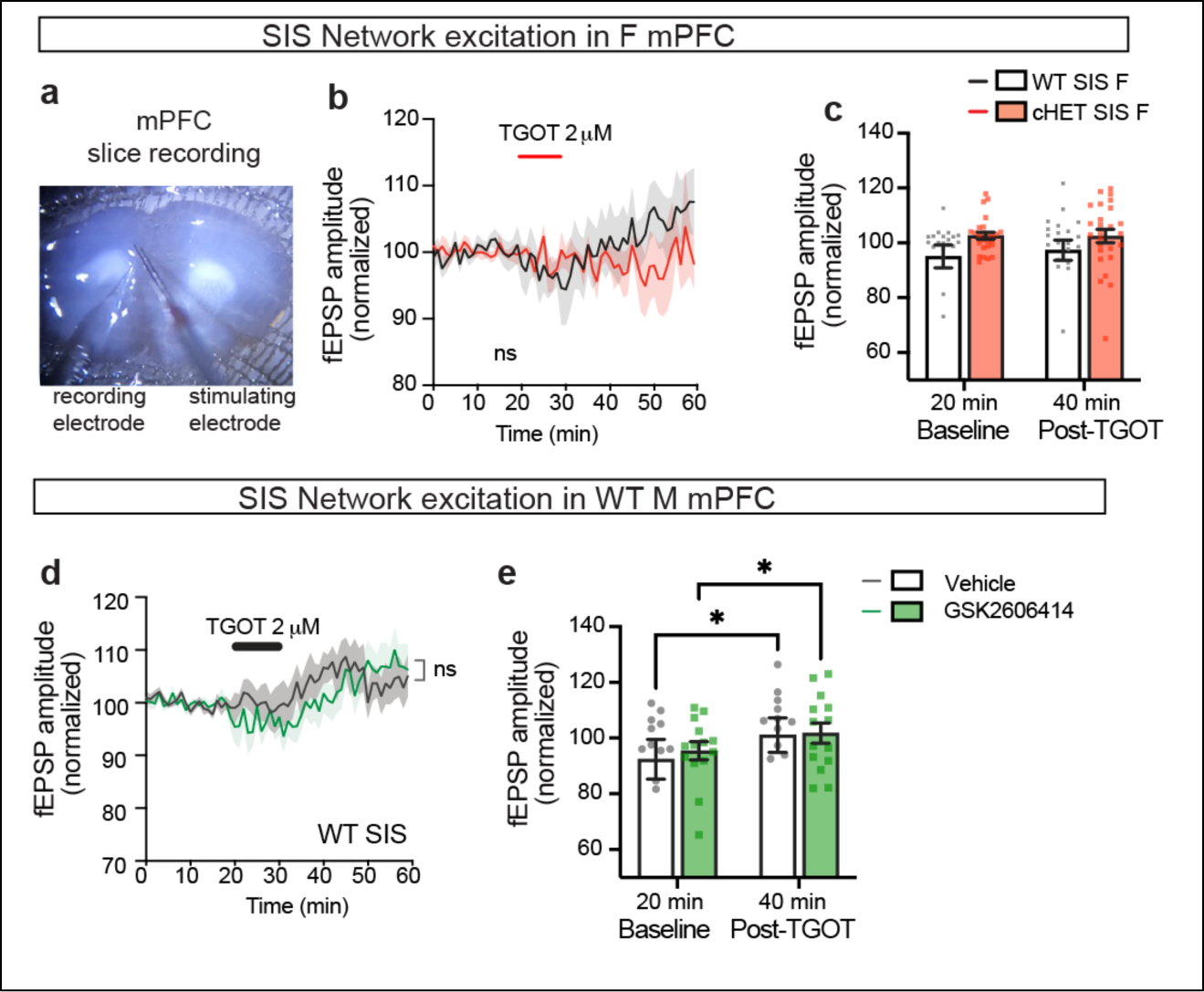
OT modulation of evoked field potentials in mPFC. **a)** Representative image of slice recording setup for mPFC. **b)** Both WT and cHET female display similar TGOT-modulation of evoked field potentials in inner layer of mPFC. **c)** No significant increase in the amplitude of field potential is observed with TGOT application for both genotypes. **c)** PERK inhibition does not alter TGOT-modulation of evoked network activity in male WT mPFC. **d)** Both vehicle and GSK2606414 treated animals have similar network excitation response to TGOT application. Statistical tests: b & d) RM Two-way ANOVA; c & e) Two-way ANOVA with Bonferroni post-hoc test. *p<0.05. n=9-14 slices from 3 animals/group.

**Extended Data Figure 10.**
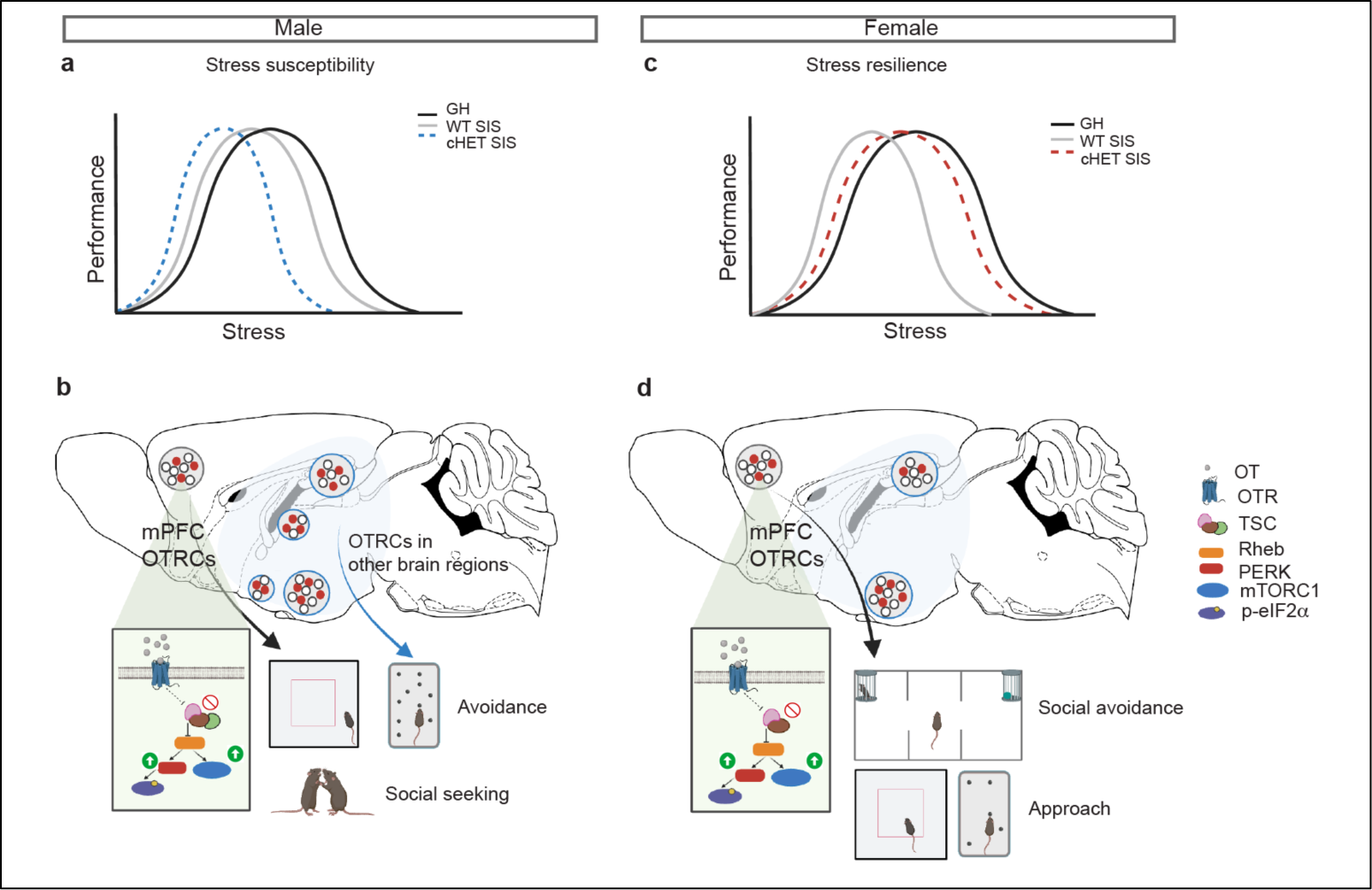
Model of stress modulation of approach-avoidance balance in male and female mice. **a)** Stress-performance U-shaped curve for male mice. Tsc2 heterozygous deletion in OTRCs amplifies the effect of social isolation stress on behavioral performance. **b)** Dysfunctional OTRCs in male mPFC contribute to maladaptive avoidance in open field and marble-burying test, and promotes social seeking for buffering the effects of stress. **c)** Stress-performance U-shaped curve for female mice. Tsc2 heterozygous deletion in OTRCs confers resilience from the effect of social isolation stress on behavioral performance. **d)** Dysfunctional OTRCs in female mPFC contribute to social avoidance but confer resilience in anxiogenic environments.

## Notes

### Competing Interest Statement

The authors have declared no competing interest.

